# Parallel genome-scale CRISPR-Cas9 screens uncouple human pluripotent stem cell identity versus fitness

**DOI:** 10.1101/2023.05.03.539283

**Authors:** Bess P. Rosen, Qing V. Li, Hyein S. Cho, Dingyu Liu, Dapeng Yang, Sarah Graff, Jielin Yan, Renhe Luo, Nipun Verma, Jeyaram R. Damodaran, Hanuman T. Kale, Samuel J. Kaplan, Michael A. Beer, Simone Sidoli, Danwei Huangfu

## Abstract

Pluripotent stem cells are defined by their self-renewal capacity, which is the ability of the stem cells to proliferate indefinitely while maintaining the pluripotent identity essential for their ability to differentiate into any somatic cell lineage. However, understanding the mechanisms that control stem cell fitness versus the pluripotent cell identity is challenging. To investigate the interplay between these two aspects of pluripotency, we performed four parallel genome-scale CRISPR-Cas9 loss-of-function screens interrogating stem cell fitness in hPSC self-renewal conditions, and the dissolution of the primed pluripotency identity during early differentiation. Comparative analyses led to the discovery of genes with distinct roles in pluripotency regulation, including mitochondrial and metabolism regulators crucial for stem cell fitness, and chromatin regulators that control pluripotent identity during early differentiation. We further discovered a core set of factors that control both stem cell fitness and pluripotent identity, including a network of chromatin factors that safeguard pluripotency. Our unbiased and systematic screening and comparative analyses disentangle two interconnected aspects of pluripotency, provide rich datasets for exploring pluripotent cell identity versus cell fitness, and offer a valuable model for categorizing gene function in broad biological contexts.

## Introduction

Self-renewal refers to the ability of stem cells to proliferate and make more stem cells. However, due to the intricate relationship between these two aspects of self-renewal, it has been challenging to distinguish the regulation of each. To tackle this fundamental question in stem cell biology, we focus on human pluripotent stem cells (hPSCs), which include embryonic stem cells (hESCs) and induced pluripotent stem cells (hiPSCs). The self-renewal of hPSCs involves mechanisms governing their pluripotent identity, ensuring their ability to differentiate into all somatic cell types, alongside mechanisms supporting their cell fitness, promoting survival and proliferation in maintenance conditions. While there are commonalities in these mechanisms, evidence also suggests the involvement of distinct regulatory pathways. The transcription factors (TFs) OCT4 (POU5F1), NANOG, and SOX2 are well-established master regulators of pluripotent identity ^1–5^ that cooperatively regulate one another as well as a large set of downstream genes in hPSCs ^6,7^. However, the downregulation of these master regulators during hPSC differentiation is accompanied by decreased susceptibility to apoptosis, with OCT4 depletion in some contexts even imparting a competitive advantage in hPSC fitness ^8,9^. In other words, pro-pluripotency factors do not always promote hPSC fitness. Consistent with this notion, large-scale loss-of-function screens that interrogate the fitness aspect of self-renewal through assaying cell survival and proliferation ^9–13^ did not rank OCT4, NANOG, and SOX2 among the top pro-fitness hits. By extension, such fitness-oriented screens may overlook other key pluripotency regulators that have weak or modest involvement in survival and proliferation. To address this need, we designed four genome-scale CRISPR-Cas9 knockout screens to separately explore genes responsible for regulating pluripotent identity and those governing stem cell fitness.

We reasoned that pluripotency in development is transient and quickly transitions into the three germ layers, thus an effective approach to unraveling the regulation of pluripotency should investigate the dissolution of pluripotent cell identity during hPSC differentiation. While previous screens have interrogated the exit from mouse naïve pluripotency ^14–16^, the distinct regulation of the mouse and human pluripotent states ^17,18^ , likely reflecting the differential regulation of naïve and primed pluripotent states ^19–21^, suggests the need for interrogation of exit from human primed pluripotency. We applied guided neuroectoderm (NE) and definitive endoderm (DE) differentiation protocols ^22,23^ to mimic the pluripotent cells in the epiblast making the first lineage commitment to differentiate into ectoderm or mesendoderm lineages at the onset of gastrulation. These lineage transitions involve the loss of the pluripotent identity and the gain of a new lineage identity. Distinct from previous hPSC differentiation screens that inspected the acquisition of a new cell fate during directed differentiation ^9,24^, we designed our pluripotency screens to examine the loss of pluripotency using a knockin fluorescent reporter for *OCT4* expression. We identified many shared hits between our NE and DE pluripotency screens, pointing to a common program orchestrating the dissolution of pluripotency independent of differentiation context. Furthermore, comparison of our pluripotency screens to previous differentiation screens ^9,24^ highlights distinct forces that drive cell state transitions: those that pull towards a new state, and those that push away from the old state.

Our pluripotency screens conducted during dynamic cell state transitions discovered genes that control pluripotent identity independent of differentiation context. We further investigated the regulation of primed human pluripotency during differentiation in comparison to results from primed hESC fitness screens ^11–13^. For better matched comparisons, we also conducted our own screens, and categorized hits into gene modules with distinct or similar impacts on pluripotent identity or cell fitness. Ontological analyses of screen hits suggest that hESC proliferation screens are more effective at uncovering genes related to stem cell fitness, such as mitochondrial components and cell cycle regulators, while our pluripotency screens are more effective at identifying regulators of the stem cell identity, many of which are chromatin factors. Defining modules containing positive regulators of both pluripotency and cell fitness, both requirements for self-renewal, allowed us to expand the core pluripotency network involving OCT4, NANOG and SOX2 through the identification of new hits such as the deubiquitinase OTUD5. Comparison of our multiple parallel CRISPR screens provides an effective gene characterization and discovery model that enabled us to disentangle two interconnected aspects of pluripotency: pluripotent cell identity versus stem cell fitness.

### Genome-scale screens for regulators of exit from pluripotency

To discover regulators of pluripotency during differentiation, we used the expression of core pluripotency and early embryogenesis regulator OCT4 ^1,2,25^ as a readout for the pluripotent state. Using our established selection free knock-in strategy ^26^, we generated an *OCT4-*GFP reporter in the H1 iCas9 hESC line, which expresses Cas9 upon doxycycline treatment ^27^ (**Supplementary data Fig. S1a,b**). *OCT4*-GFP decreased upon NE and DE differentiation (**Supplementary data Fig. S1c,d**), mirroring the downregulation of *OCT4* expression during the dissolution of the pluripotency network. We next conducted parallel screens in NE and DE differentiation conditions to distinguish whether the regulation of pluripotency was largely differentiation context dependent, or if a shared group of regulators might control the dissolution of pluripotency regardless of differentiation context. Using a pooled screening strategy ^24,28,29^, we infected H1 OCT4^GFP/+^ iCas9 hESCs with a genome-scale CRISPR library ^30^ (**Fig. 1a**). To investigate the dissolution of pluripotent identity during differentiation, library infected H1 iCas9 *OCT4^GFP/+^* hESCs were differentiated in NE and DE differentiation conditions. OCT4-GFP^hi^ and OCT4-GFP^lo^ were isolated by fluorescence-activated cell sorting (FACS) on day 1.5 of NE (**Supplementary data Fig. S2a,b**) and day 2.5 of DE (**Supplementary data Fig. S2c,d**) differentiations, matching the timepoints when *OCT4*-GFP was rapidly downregulated (**Supplementary data Fig. S1c**). The MAGeCK robust ranking aggregation (RRA) algorithm ^31^ was used to identify genes that promote or inhibit pluripotency based on relative gRNA enrichment in the OCT4-GFP^lo^ or OCT4-GFP^hi^ population, respectively. Using a false discovery rate (FDR) cutoff of 0.1 we identified 90 pro-pluripotency and 72 anti-pluripotency hits in NE, and 33 pro-pluripotency and 65 anti-pluripotency hits in the DE screen (**Supplementary Table 1**). These results suggest that the NE screen condition was more robust than the DE screen condition for identifying positive regulators of pluripotency, which may explain why *OCT4* was identified as a top pro-pluripotency hit in the NE but not DE screen.

**Figure 1.**
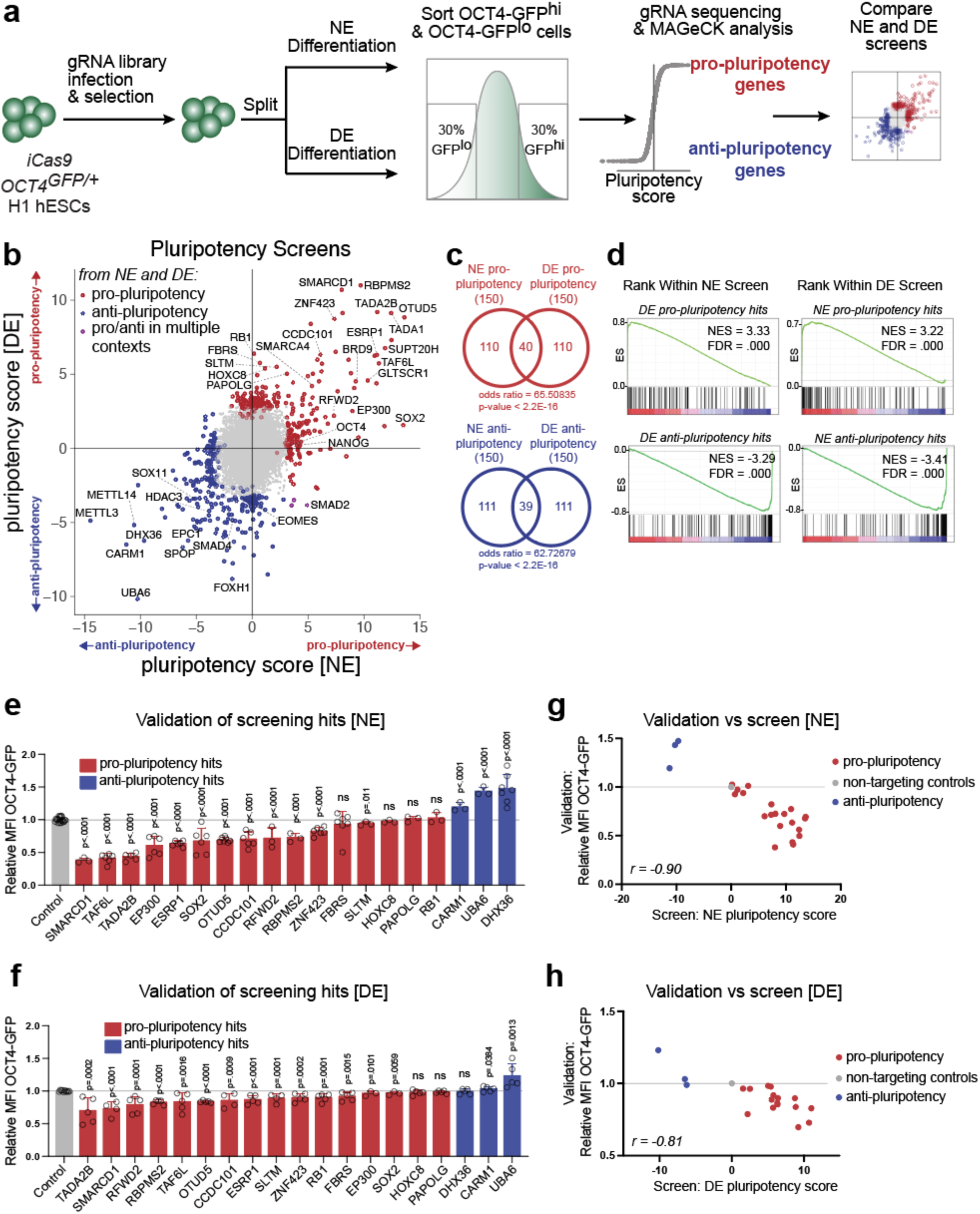
NE and DE screens for exit of pluripotency reveal common program. (a) Schematic of NE and DE CRISPR screen comparison. (b) NE and DE Screen results by pluripotency score per gene = log10(RRA score OCT4-GFP^hi^ enrichment) − log10(RRA score OCT4-GFP^lo^ enrichment), top 150 pro-pluripotency (ranked by RRA score OCT4-GFP^lo^ enrichment) and top 150 anti-pluripotency (ranked by RRA score OCT4-GFP^hi^ enrichment) hits from each screen labelled, RRA scores determined by MAGeCK. (c) Overlap of top 150 pro-pluripotency genes identified in NE vs. DE context screens, and top 150 anti-pluripotency genes identified in NE vs. DE context screens, Fisher’s exact test used for comparison. (d) GSEA for enrichment of pro- and anti-pluripotency top hits in screen results ranked by pluripotency score, NE vs. DE screens. (e) Bar graphs show relative MFI of OCT4-GFP following gRNA targeting of H1 iCas9 *OCT4^GFP/+^* hESCs during NE differentiation. Relative MFI OCT4-GFP = (MFI per gRNA) / (MFI non-targeting controls). *n*=3-6 independent experiments. 2 non-targeting controls analyzed per experiment. Data represented as mean. Error bars indicate s.d. Statistical analysis was performed by unpaired two-tailed Student’s *t*-test. P-values indicated. (f) Bar graphs show relative MFI of OCT4-GFP following gRNA targeting of H1 iCas9 *OCT4^GFP/+^* hESCs during DE differentiation. Relative MFI OCT4-GFP = (MFI per gRNA/ MFI in-well tdT uninfected control) / (MFI non-targeting controls/ MFI in-well tdT uninfected control). *n=* 5 independent experiments. 2 non-targeting controls analyzed per experiment. Data represented as mean. Error bars indicate s.d. Statistical analysis was performed by unpaired two-tailed Student’s *t*-test. P-values indicated. (g) Comparison NE screen pluripotency score vs. NE validation mean relative intensity OCT4-GFP by gene. Statistical analysis by Pearson correlation test. (h) Comparison DE screen pluripotency score vs. DE validation mean relative intensity OCT4-GFP by gene. Statistical analysis by Pearson correlation test.

To allow for comparison between screens performed in different contexts, we selected an inclusive top 150 genes in each category as hits for further analysis based on RRA scores. To assist comparisons between the NE and DE screens, we also calculated a “pluripotency score” for each gene in each screen equal to log10(RRA score OCT4-GFP^hi^ enrichment) – log10(RRA score OCT4-GFP^lo^ enrichment) (**Supplementary Table 1**). A positive pluripotency score indicated a gene was a positive regulator of pluripotent identity (pro-pluripotency): knockout hastens the loss of pluripotent identity during differentiation. A negative pluripotency score indicated a gene was a negative regulator of pluripotent identity (anti-pluripotency): knockout slows the loss of pluripotent identity during differentiation. Despite the screens being performed in different differentiation contexts, the results of the NE and DE screens were moderately correlated (Pearson correlation coefficient *r =* 0.56), with a substantial number of overlapping hits: 40 hits that promote pluripotency (pro-pluripotency) and 39 hits that inhibit pluripotency (anti-pluripotency) (**Fig. 1b, c**). Gene set enrichment analysis (GSEA) revealed significant enrichment (Normalized Enrichment Score, NES, greater than 3.2) of pro-pluripotency hits from one screen in genes ranked by the pluripotency scores in the other screen; and strong negative enrichment (NES less than –3.2) of anti-pluripotency hits (**Fig. 1d**). The high degree of concordance between NE and DE screens suggests the presence of common mechanisms controlling the loss of pluripotency regardless of the external differentiation cues.

Some of the anti-pluripotency hits identified in our screens are known to promote differentiation. For instance, *SOX11* is a neural regulator ^32^, and *FOXH1*, *SMAD2*, *SMAD4*, and *EOMES* regulate DE specification in mice and humans ^24,33–36^. Therefore, we compared pro- and anti-pluripotency hits found in our screens to the anti- and pro-differentiation hits identified from previous screens interrogating the specification of SOX17^+^ DE and EPCAM^-^/NCAM^+^ NE fates ^9,24^ (**Supplementary data Fig. S3a,b**). To allow comparison between multi-context screens performed using varying protocols, hits were defined by top 150 genes by previously published RRA score (**Supplementary Table 2**). A small number of genes were identified as both pro-pluripotency and anti-differentiation, or both anti-pluripotency and pro-differentiation, supporting the interconnectivity of the acquisition of the new cell identity and the dissolution of the pluripotent state (**Supplementary data Fig. S3c,d**). However, most of the pluripotency hits were not identified in previous differentiation screens. For a better matched comparison, we compared the DE pluripotency screen with our previous DE differentiation screen ^24^ conducted under closely matched differentiation and screening conditions, and conducted systemic analysis of overlapping and non-overlapping hits using STRING (v11.5) ^37^ and GO term enrichment ^38^ (**Supplementary data Fig. S3e,f**). The results showed that top hits unique to the DE differentiation screen included regulators of developmental signaling pathways, and Golgi-associated transport and processing. In contrast, top hits unique to the pluripotency screen were chromatin modifiers and transcriptional regulators, and genes involved in RNA processing. The different hits identified from screens performed in the same differentiation context suggests that the gene network regulating the pull of differentiation can be decoupled from the push of pluripotency loss.

With the aim of identifying specific regulators of pluripotency, rather than differentiation, we selected 19 genes not known to be regulators of lineage specification for further validation: 16 genes identified as pro-pluripotency in at least one screen, and 3 genes identified as anti-pluripotency. Hits were chosen for validation by RRA score with an effort to represent 4 different hit types during validation: identified as pro-pluripotency in NE and DE screens, identified as pro-pluripotency in either NE or DE screens, and identified as anti-pluripotency (**Supplementary Table 2**). Lentiviral vectors expressing gene-specific gRNAs were used to infect H1 OCT4^GFP/+^ iCas9 hESCs followed by individual differentiation assays. NE differentiation for validation was performed as in original screening conditions (**Extended Data Figure 4A**). For DE differentiation, which exhibits a high sensitivity to variations in cell density across experiments, we compared gRNA-infected hPSCs with the control *tdTomato* infected H1 OCT4^GFP/+^ iCas9 line, co-cultured in the same well (see Methods). Using *OCT4*-GFP mean fluorescent intensity (MFI) relative to non-targeting controls as measured by flow cytometry, we validated 15 and 16 hits in NE and DE differentiation, respectively (**Fig. 1e,f, Supplementary data Fig. S4a-d**). Validation phenotypes correlated strongly with pluripotency scores in each screen (**Fig. 1g,h**). Hits validated in both NE and DE contexts included genes not previously connected to the regulation of pluripotency, such as *OTUD5*, *ZNF423*, *SLTM,* and *UBA6*. Validation results in NE and DE contexts were also well correlated (**Supplementary data Fig. S4e**), and several genes (e.g., *RFWD2*, *SOX2*, *EP300*, and *SLTM*) identified among the top 150 hits in only one screen were validated in both NE and DE contexts (**Fig. 1e-f, Supplementary Table 2**), despite the less robust nature of the DE differentiation system as an assay for pluripotency. Our screening and validation studies reveal the existence of a common program that controls the dissolution of pluripotency across different differentiation contexts, and this program is distinct from those that govern lineage specification.

### Parallel Screens reveal distinct regulation of pluripotent identity and fitness

To compare the regulation of pluripotent identity in differentiation contexts to regulation of cell fitness in maintenance conditions, we conducted two additional pooled genome-scale CRISPR knock-out screens measuring hESC proliferative capacity. The first screen was performed in the chemically defined serum-free E8 media ^39^ containing FGF2 and TGFβ, both essential for primed hESC maintenance (**Fig. 2a, Supplementary data Fig. S5a**); and the second screen used a “challenge” condition wherein cells were exposed to E6 media (similar to E8 but without FGF2 and TGFβ) for five days before returning the cells back to the standard E8 condition for one passage (**Fig. 2a, Supplementary data Fig. S5b**). We used the MAGeCK RRA algorithm ^31^ to identify genes that promote or inhibit fitness based on relative gRNA enrichment in Day 0 or Day 10, respectively. We evaluated the screen quality by calculating precision-recall curves based on previously defined essential and non-essential gene sets derived from 78 CRISPR knockout screens ^40^. Both the E8 and E6 screens had Area Under the Curve (AUC) scores exceeding 0.91. In contrast, the NE and DE screens, which do not evaluate essentiality, exhibited poor performance as expected (**Supplementary data Fig. S5e**).

**Figure 2.**
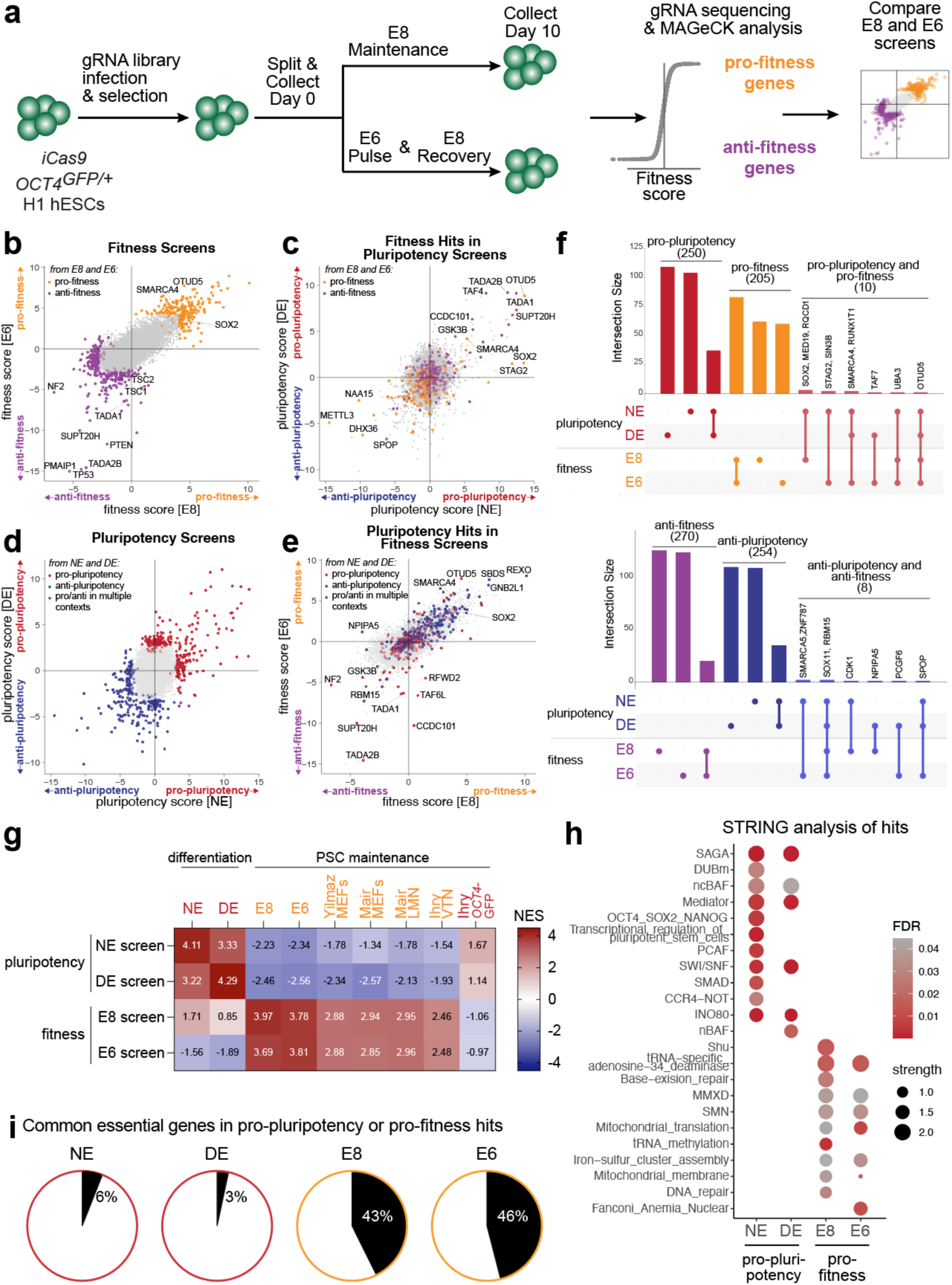
Comparison pluripotency and cell fitness screens reveal distinct hits. (a) Schematic E8 and E6 fitness screens. (b) E8 and E6 Screen results by fitness score per gene = log10(RRA score Day 10 enrichment) − log10(RRA score Day 0 enrichment), top 150 pro-fitness (ranked by RRA score Day 0 enrichment) and top 150 anti-fitness (ranked by RRA score Day 10 enrichment) hits from each screen labelled. (c) E8 and E6 screen fitness hits in NE and DE screens. (d) NE and DE Screen results by pluripotency score per gene = log10(RRA score OCT4-GFP^hi^ enrichment) − log10(RRA score OCT4-GFP^lo^ enrichment), top 150 pro-pluripotency (ranked by RRA score OCT4-GFP^lo^ enrichment) and top 150 anti-pluripotency (ranked by RRA score OCT4-GFP^hi^ enrichment). (e) NE and DE screen pluripotency hits in E8 and E6 screens. (f) Upset plot ^95^ of overlap NE, DE pro-pluripotency hits and E8, E6 pro-fitness, and Upset plot overlap NE, DE anti-pluripotency and E8, E6 anti-fitness hits. (g) GSEA for enrichment of pro-pluripotency and pro-fitness gene sets from NE, DE, E8, E6 screens, previous screens by Yilmaz et al. ^11^, Mair et al. ^12^, and Ihry et al. ^13^, as re-analyzed by Mair et al. , and previous pluripotency screen as published in Ihry et al. ^13^. Gene sets from previous fitness screens are top 150 genes per screen as ranked by Bayes factor score, calculated by BAGEL analysis ^97^. Gene sets from previous pluripotency screen are top 150 genes per screen as ranked by RSA score^98^. Pluripotency screens labelled in red text, fitness screens labelled in orange text. (h) STRING^37^ database analysis pro-pluripotency and pro-fitness hits from NE, DE, E8, and E6 screens, strength = log10(observed/expected) for enrichment of gene types within a set. (i) Proportion of common essentiality genes as defined by DepMap data sets ^51,53,87^ identified as top pro-pluripotency or pro-fitness hits in NE, DE, E8, and E6 screens.

Using an FDR cutoff of 0.1, we identified 323 pro-fitness and only 1 anti-fitness hit in the E8 screen, and 357 pro-fitness and 46 anti-fitness hits in the E6 screen. Given that the E8 condition is already optimized for proliferation, the E6 challenge condition is more conducive to the identification of anti-fitness hits. To allow for comparison between screens performed in different contexts, we selected an inclusive top 150 genes in each category as hits for further analysis based on RRA scores. To assist comparisons between the E8 and E6 screens, we also calculated a “fitness score” for each gene equal to log10(RRA score Day 10 enrichment) – log10(RRA score Day 0 enrichment) (**Supplementary Table 1**). There were large numbers of overlapping pro-fitness (85) between the E8 and E6 screens (**Fig. 2b, Supplementary data Fig. S5c**), and GSEA analysis revealed strong enrichment of hits identified from one screen in genes ranked by the fitness scores in the other screen (**Supplementary data Fig. S5d**). Some shared anti-fitness (24) hits were also identified, including known negative regulators of hESC self-renewal such as *TP53* and *PMAIP1.* The E6 screen also identified additional hits, including *PTEN*, *TSC1*, and *TSC2*, which were identified in a previous CRISPR screen ^11^.

We assessed the degrees of correlation between hits in all four screens based on fitness and pluripotency scores. Hits from the E8 and E6 screens showed a strong correlation in self-renewal scores (Pearson correlation coefficient *r =* 0.82), and hits from the DE and NE screens also showed correlated pluripotency scores as described. In contrast, we did not observe any positive correlation between fitness and pluripotency scores across the two screening types (**Supplementary data Fig. S5f**). Consistent with the correlation analysis, we found that fitness screening hits were not highly ranked in the pluripotency screens (**Fig. 2c**), while pluripotency screen hits (**Fig. 2d**) were not highly ranked in the fitness screens (**Fig. 2e**). Indeed, very few genes were identified as promoting both pluripotency and fitness or, conversely, inhibiting both pluripotency and fitness (**Fig. 2f**). We expanded the comparison to include the top 150 pro-fitness hits from previous fitness screens in a variety of hESC maintenance conditions ^11–13^ (**Supplementary Table 3**) as gene sets for GSEA analysis. As expected, there was a high degree of enrichment of hits from these previous fitness screens in the E8 and E6 screen results (NES ranging from 2.46 to 3.97) but not in the DE and NE pluripotency screen results (**Fig. 2g, Supplementary data Fig. S5g**). However, when utilizing the top 150 pro-pluripotency hits from a screen based on the expression of OCT4-GFP in hESC maintenance conditions (referred to as the Ihry OCT4-GFP screen) ^13^, we observed a different pattern. GSEA analyses revealed a lack of enrichment in our E8 and E6 screens, but a modest enrichment in NE and DE pluripotency screens. These results suggest that screens focusing on cell identity, based on OCT4 expression, exhibit greater similarity to each other and are distinct from screens focusing on cell fitness. Indeed, shared hits between the Ihry OCT4-GFP screen and our NE/DE pluripotency screens included well-known pluripotency regulators such as OCT4 and NANOG (**Supplementary data Fig. S5h**), though the NE and DE screens assaying the exit of pluripotency shared a larger number of hits with one another than with the Ihry OCT4-GFP maintenance screen.

Analysis with STRING ^37^ showed distinct protein complexes were discovered in pluripotency and fitness top hits (**Fig. 2h**). The NE and DE screens identified many chromatin regulators. Some of these have been implicated in pluripotency regulation such as the Spt-ADA-Gcn5-Acetyltransferase (SAGA) complex ^41,42^, the transcriptional regulating mediator complex ^43^, the chromatin remodeling INO80 complex ^44^, and the ATP dependent nucleosome remodeling BRG1-associated factor (BAF) complex (also known as the SWI/SNF complex) ^21,45–47^. In contrast, complexes identified as pro-fitness in the E8 and E6 screens included mitochondrial components and regulators of DNA-repair and tRNA metabolism, consistent with the findings of previous fitness screens ^11,12^. We performed Reactome pathway analysis ^48^ on hits from each screen, then ranked sets by median absolute log_2_(fold change of screening enrichment) (LFC) of all hits in set (**Supplementary data Fig. S6a**). Members of the “chromatin organization”, “HATs acetylate histones”, and related Reactome sets had a high median |LFC| in NE and DE pluripotency screens, whereas genes from the “mitochondrial translation”, “tRNA aminoacylation”, and related sets had a high median |LFC| in E8 and E6 fitness screens. The overrepresentation of mitochondrial components and tRNA processing genes in pro-fitness hits, as revealed by both STRING and Reactome analyses, confirm that the fitness screens preferentially capture genes required for cell proliferation and survival, rather than cell identity. Supporting this notion, >40% of pro-fitness hits in our E8 and E6 screens are categorized as common essential genes based on integrated screening results from hundreds of cancer cell lines ^49–53^ (**Fig. 2i, Supplementary data Fig. S6b, Supplementary Table 3**). Similarly, a high proportion of common essential genes was also found in previous hESC fitness screens ^11–13^ (**Supplementary data Fig. S6c**). In contrast, fewer than 10 pro-pluripotency hits from the NE or DE screens overlapped with common essential genes. Together, these results indicate that the fitness screens predominantly identify regulators of stem cell fitness, while our pluripotency screens in differentiation contexts mainly uncover regulators of the stem cell identity. In addition, chromatin modulation plays a key role in regulating pluripotency during cell fate transition.

### Comparative analysis of screening results defines gene modules

To dissect gene modules with distinct roles regulating pluripotency and hESC fitness, we performed unbiased hierarchical clustering of all pro-pluripotency and pro-fitness hits by the column scaled LFC in all 4 screens (**Fig. 3a, Supplementary Table 4**). This approach revealed 8 gene modules that exhibited distinct patterns in their pluripotency and fitness scores (**Fig. 3b**). Reassuringly, analysis of individual modules by STRING showed association between proteins corresponding to genes in the same module, suggesting that our clustering method based on findings from genetic screening data is effective for grouping hits with similar biochemical functions and localizations (**Supplementary data Fig. S7**). Many of these modules were primarily enriched for either pro-pluripotency or pro-fitness hits, and reflected broader screen findings regarding the regulation of pluripotent identity and hESC proliferation and survival. For example, module 1, 4 and 5, which consisted largely of pro-fitness hits, were enriched for regulators of mitochondrial translation, mitochondrial respiration, or both. This is consistent with our and previous findings connecting mitochondrial regulation with essentiality in pluripotency ^11^. In comparison, module 3 and 2, both containing pro-pluripotency hits, were enriched for genes involved in the signaling pathways regulating pluripotency including WNT, TGFβ, and Hippo signaling, and a host of developmental TFs, respectively. Module 6 was notable for its enrichment of genes that were identified as strongly pro-pluripotency but anti-fitness, which included many members of the SAGA histone acetyltransferase and core complex, the mediator complex, and the recently identified non-canonical (ncBAF) complex ^54,55^ (**Fig. 3c**). Overall, STRING analyses revealed great interconnectivity between components within each module. The gene relationships identified in these modules demonstrate both the utility of clustering multi-context screening data to identify distinct regulatory programs, and the potential for inferring the biochemical functions of uncharacterized genes.

**Figure 3.**
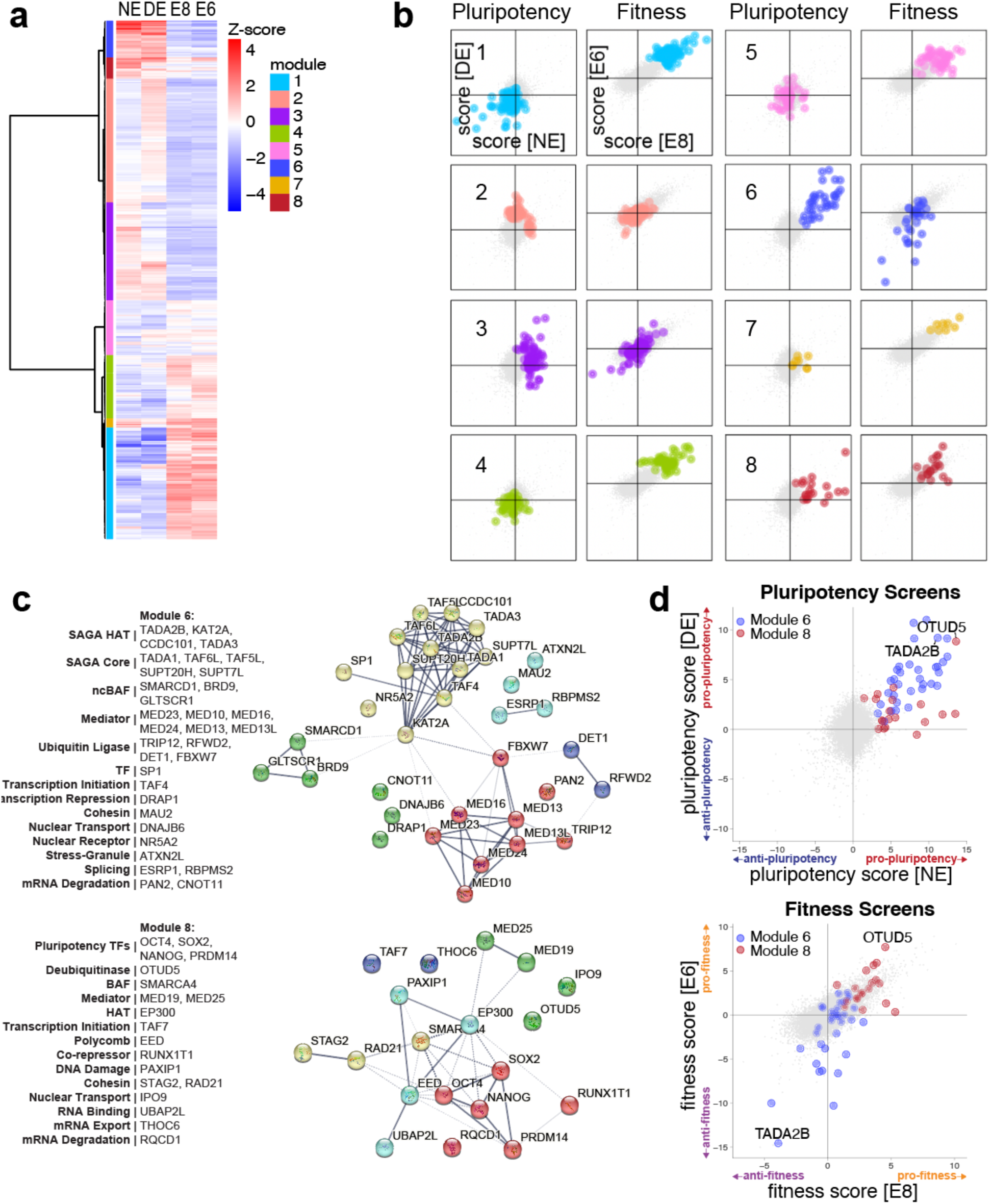
Defining gene modules based on screening results. (a) Hierarchical clustering by z-score-per-gene of pro-pluripotency hits from NE and DE screens, and pro-fitness hits from E8 and E6 screens. Relative levels of LFC represented as column z-score. (b) Modules by pluripotency score in NE and DE screens, and fitness score in E8 and E6 screens. (c) Annotation of all genes within modules 6 and 8, and Interactome indicating known associations between members of module gene sets identified through STRING^37^ analysis using default parameters for high-confidence interactions. Each gene-set was further clustered using k-means clustering (k=5), and dotted lines indicate connections between clusters. (d) Modules 6 and 8 in NE and DE screens by pluripotency score, and E8 and E6 screens by fitness score with archetypal genes *TADA2B* and *OTUD5* labelled.

One module, module 8, was noteworthy in that it was highly enriched for genes that were highly ranked as both pro-pluripotency (in NE, DE, or both) and pro-fitness (in E8, E6 or both). Module 8 included all 3 core pluripotency factors OCT4, NANOG, and SOX2, as well as PRDM14, a TF which is essential to the maintenance of human pluripotency ^44,56,57^ (**Fig. 3c**). There were also chromatin remodelers associated with pluripotency including coactivator EP300 ^58^, Polycomb complex member EED ^59^, mediator complex members MED19 and MED25 ^43^, and BAF ATPase SMARCA4 ^21,45–47,54,55,60^. Notably, analysis by STRING showed a great degree of functional protein association within this module (**Fig. 3c**). The large number of module 8 members involved with chromatin accessibility and transcriptional control of pluripotency suggests a similar role for other hits in this module such as OTUD5 and RUNX1T1 that were not previously connected to pluripotency.

### OTUD5 and TADA2B are pro-pluripotency genes with opposing roles in cell fitness

Module 6 and module 8 from our comparative analysis both captured genes with high pluripotency scores, yet they were differentiated by enrichment with genes with opposing effects on cell fitness (**Fig. 3b,c**). For further investigation, we chose two archetypical examples of screen hits from each of these modules: *OTUD5* (in module 8) and *TADA2B* (in module 6) (**Fig. 3d**). Both genes were already validated for their pro-pluripotency roles (**Fig. 4e,f**), so we set out to confirm their opposing roles in hESC fitness. We used CRISPR-Cas9 targeting to generate multiple clonal knockout (KO) lines of *TADA2B* and *OTUD5*, which were confirmed by sequencing and western (**Fig. 4a-d**). Consistent with our screening results, *TADA2B* KOs demonstrated a higher growth rate than WT in E8 and E6 conditions (**Fig. 4e,f**), which is consistent with previous findings ^9^. In contrast, *OTUD5* KOs displayed reduced growth compared to WT in E6 medium (**Fig. 4h**). However, *OTUD5* KO hESCs did not display a significant growth defect in the standard hESC E8 culture condition (**Fig. 4g**). To reconcile this finding with the E8 screening results, we speculate that *OTUD5* KO hESCs have compromised cell fitness, but the phenotype could be masked under optimal culture conditions. Supporting this idea, we found that *OTUD5* KO hESCs showed reduced growth in the E8 condition when the cells were challenged by passage without the Rho-associated protein kinase (ROCK) inhibitor Y-27632 (**Fig. 4i**), a compound normally used for optimal survival of dissociated hESCs during passaging. We further speculate that hESCs were more vulnerable in a large-scale CRISPR screening setting not only because the culture condition may be less-than-ideal, but also because of cell-cell competition. To test the latter idea, we used lentiviruses to barcode individual KO and WT lines and cultured the cells together for a period of four weeks while collecting cells weekly to assess the representation of barcoded lines in culture. Representation of *OTUD5* KOs were greatly reduced over time while the representation of WT cells increased (**Fig. 4j**). The divergence in fitness phenotype between module 6 and module 8, confirmed by phenotypic analysis of archetypal gene knockouts, indicates that pro-pluripotency genes may exert opposing effect on cell fitness. These findings underscore the complexity of the relationship between cell fitness and pluripotent identity. As such, it is not sufficient to use fitness in hESC growth conditions as a proxy for pluripotent identity.

**Figure 4.**
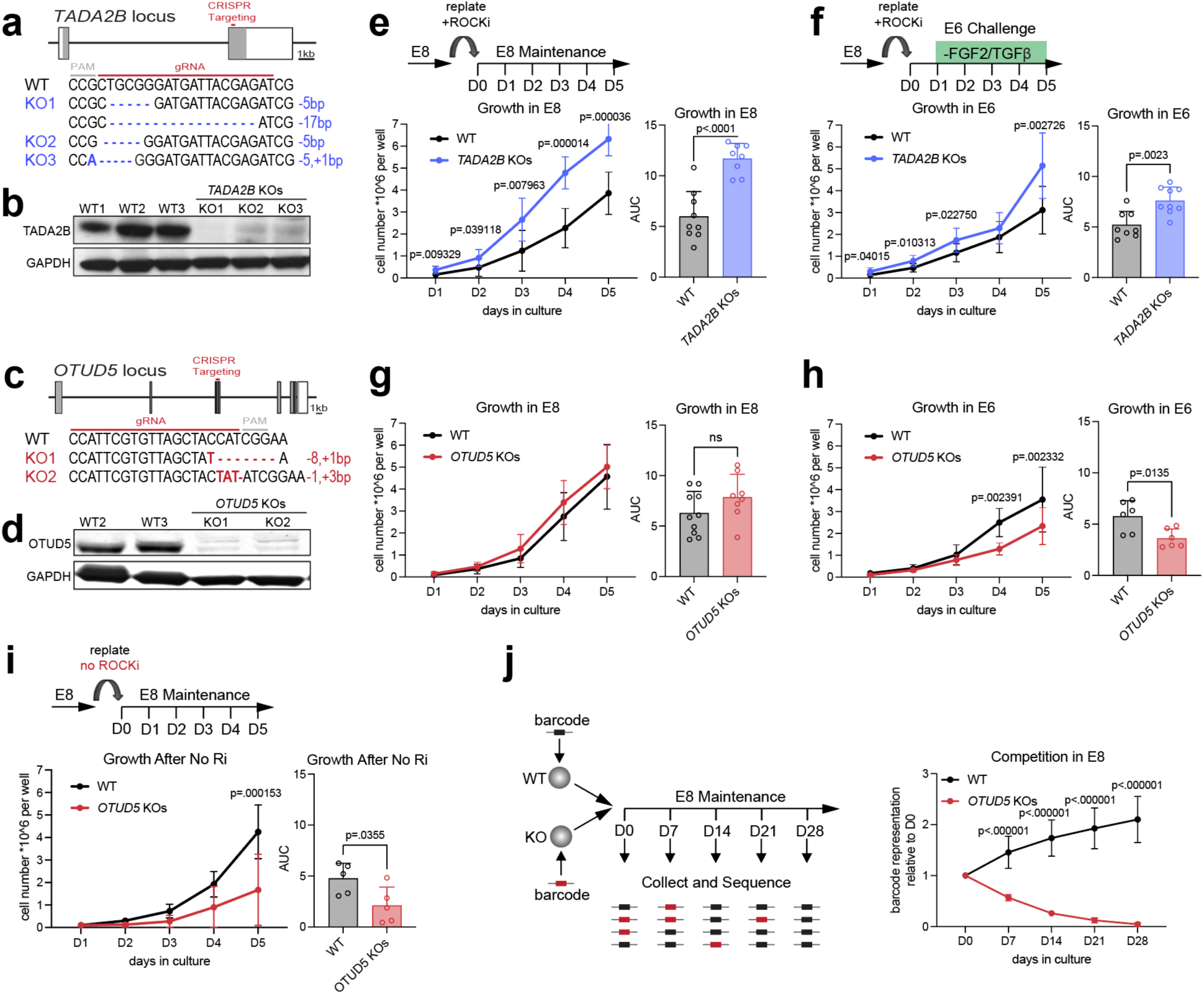
*TADA2B* KOs and *OTUD5* KOs have opposing fitness phenotype. (a) Schematic of *TADA2B* including location of gRNA targeting and sequences of knockouts (KOs). Boxes indicate *TADA2B* exons, and filled grey boxes indicate the coding sequence of *TADA2B*. (b) Western blot of TADA2B confirming KOs in HUES8iCas9 WT1, WT2, WT3, and HUES8iCas9 *TADA2B^-/-^* KO1, KO2, and KO3 clonal lines. (c) Schematic of *OTUD5* targeting and KOs. (d) Western blot of OTUD5 confirming KOs in HUES8iCas9 WT2, WT3, and HUES8iCas9 *OTUD5^-/-^* KO1, and KO2 clonal lines. (e) Growth curve and area under growth curve (AUC) of WT and *TADA2B* KOs in E8 maintenance conditions and (f) E6 challenge conditions. 3 WT and 3 KO lines analyzed per experiment. *n=* 3 independent experiments. Data represented as mean. Error bars indicate s.d. Statistical analysis was performed by unpaired two-tailed Student’s t-test. P-values for significant comparisons indicated. (g) Growth curve and AUC of WT and *OTUD5* KOs in E8 maintenance conditions. 2 WT and 2 KO lines analyzed per experiment. *n=* 5 independent experiments. Data represented as mean. Error bars indicate s.d. Statistical analysis was performed by unpaired two-tailed Student’s t-test. No significant comparisons were found. (h) Growth curve and AUC of WT and *OTUD5* KOs in E6 challenge conditions. 2 WT and 2 KO lines analyzed per experiment. *n=* 3 independent experiments. Data represented as mean. Error bars indicate s.d. Statistical analysis was performed by unpaired two-tailed Student’s t-test. P-values for significant comparisons indicated. (i) Growth curve and AUC of WT and *OTUD5* KOs after plating without Y-27632 (ROCKi), a standard component of hESC culture. 2 WT and 2 KO lines analyzed per experiment. *n=* 3 independent experiments. Data represented as mean. Error bars indicate s.d. Statistical analysis was performed by unpaired two-tailed Student’s t-test. P-values for significant comparisons indicated. (j) Cell competition assay WT vs. *OTUD5* KOs. Individual cell lines were labelled with LARRY barcodes ^105^, and pooled. Pooled cells were expanded in E8 conditions for 28 days. Pools were collected and sequenced for representation of barcodes every 7 days. WT2 and KO1, KO2 lines analyzed per experiment. *n=* 5 independent experiments. Data represented as mean. Error bars indicate s.d. Statistical analysis was performed by unpaired two-tailed Student’s t-test. P-values for significant comparisons indicated. ^38^

### TADA2B and SAGA Complex Knockouts illustrate distinct regulation of pluripotency and cell fitness

Our discovery regarding the dual role of *TADA2B*, exhibiting both anti-fitness and pro-pluripotency characteristics, may initially seem paradoxical but aligns with previous findings. *TADA2B* was identified as a top anti-fitness hit in screens of pluripotent cell proliferation, across multiple growth conditions ^9,61^. On the other hand, the SAGA complex is pro-pluripotency in mPSCs ^41,42^ and knockout of SAGA complex members, including *TADA2B*, reduces OCT4 expression in hESCs ^9^. To address this apparent paradox, we set up to better understand the mechanism by which *TADA2B* KOs enhance cell fitness: we measured markers of cell death and proliferation in WT vs. *TADA2B* KO hESCs. We detected a significantly decreased number of apoptotic markers cleaved caspase-3 (**Fig. 5a**) and Annexin-V (**Fig. 5b**) positive cells in *TADA2B* KOs. While mitotic marker phosphohistone H3, associated with proliferation, was unchanged in *TADA2B* KOs (**Fig. 5c**). This suggests that the enhanced fitness of *TADA2B* KO hESCs is primarily driven by reduced cell death rather than enhanced growth. Apoptosis-mediated cell death has a protective role in hESCs, eliminating abnormal and DNA-damaged stem cells from the population ^8,62,63^. We hypothesized that because of this reduction in cell-death *TADA2B* KOs may be unable to eliminate differentiating cells from the hESC population. To test this hypothesis, we conducted GSEA analysis on RNA-seq data comparing *TADA2B* KO with WT hESCs. We observed a negative enrichment of pluripotency-related gene sets ^64,65^ and a positive enrichment for differentiation-related gene sets (**Fig. 5d-f**). These results suggest that loss of *TADA2B* promotes survival at the expense of pluripotency. Though initially counterintuitive, these *TADA2B* KO results demonstrate the distinct regulation of two characteristics essential for hPSC self-renewal: maintenance of pluripotent identity, and robust cell survival and proliferation.

**Figure 5.**
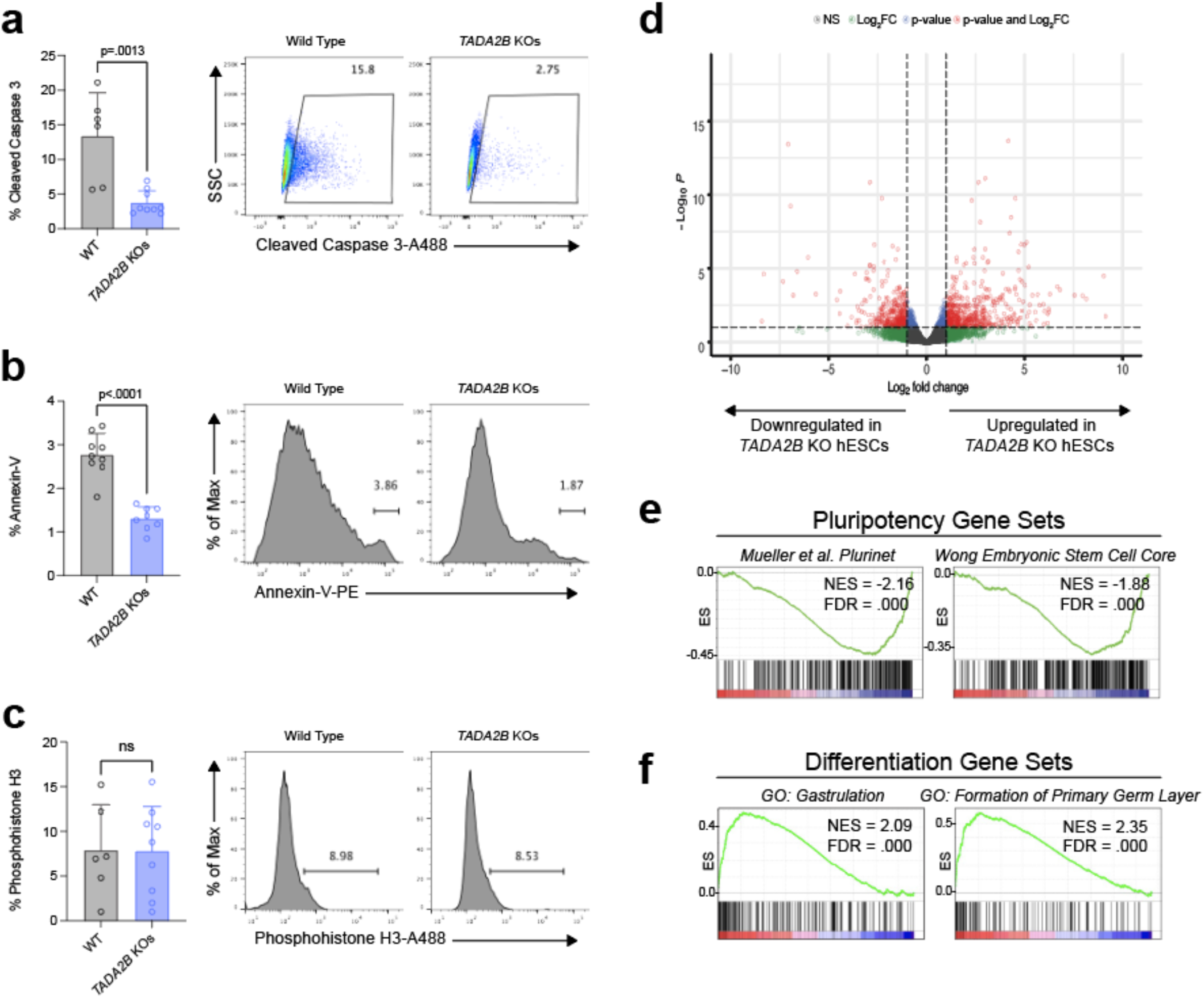
*TADA2B* KOs display reduced markers of apoptosis and increased markers of differentiation. (a) Flow cytometry quantification and representative dot-plots cleaved caspase 3 in WT and *TADA2B* KO hESCs. HUES8iCas9 WT1, WT2, WT3, and HUES8iCas9 *TADA2B^-/-^* KO1, KO2, and KO3 clonal lines analyzed in each experiment. *n=* 3 independent experiments. Error bars indicate s.d. Statistical analysis was performed by unpaired two-tailed Student’s t-test. P-values indicated. (b) Flow cytometry quantification and representative histograms Annexin-V WT and *TADA2B* KO hESCs. *n=* 3 independent experiments. (c) Flow cytometry quantification and representative histograms. Phosphohistone H3 WT and *TADA2B* KO hESCs. *n=* 3 independent experiments. (d) Volcano plot showing differential expression in *TADA2B* KO vs. WT hESCs. 3 WT and 3 KO lines analyzed by RNA-Seq. Statistical analysis with DeSeq2. (e) GSEA shows negative enrichment of pluripotency markers ^64,65^, and (f) positive enrichment of early differentiation gene sets in *TADA2B* KO vs. WT differential expression.

*TADA2B* was identified in module 6, which was particularly enriched with other components of the histone acetyltransferase and core components of the SAGA complex; in contrast SAGA complex deubiquitinase components were identified in module 3 (**Supplementary data Fig. S8a**), suggesting that SAGA core components or histone-acetyltransferase components may have a similar role in pluripotency to *TADA2B*. To investigate the role of additional SAGA complex members identified in module 6 by comparative screen analysis, lentiviral vectors expressing gene-specific gRNAs were used to infect H1 OCT4^GFP/+^ iCas9 hESCs followed by expansion in E8 or E6 conditions (**Supplementary data Fig. S8b,c**). Followed by measurement of *OCT4*-GFP mean fluorescent intensity (MFI) relative to non-targeting controls as measured by flow cytometry and cell counts. The knockout of SAGA complex members *TAF5L, TAF6L, TADA1,* and *CCDC101* in addition to *TADA2B* was shown to reduce *OCT4-*GFP expression during hESC maintenance (E8) conditions (**Supplementary data Fig. S8d**). At the same time, increased cell numbers were observed in E6 challenge conditions (**Supplementary data Fig. S8e**), suggesting that other SAGA complex members identified in module 6 play a similar role to *TADA2B* in the regulation of self-renewal: their loss resulting in a gain in cell fitness at the expense of pluripotent identity. While the reduction in *OCT4*-GFP levels with SAGA component knockout was identified in NE and DE differentiation screens, these results in E8 and E6 conditions demonstrate that multiple members of the SAGA complex are vital to the maintenance of the pluripotent state in hPSCs, which would not have been identified solely using fitness screens. Further, the reduction in pluripotency markers in SAGA complex knockouts with high cell counts emphasizes the distinction between the two interlinked aspects of hESC self-renewal: pluripotent identity and cell fitness.

### *OTUD5* knockouts confirm role of OTUD5 in the dissolution of pluripotency

In contrast to *TADA2B,* top-ranked pro-pluripotency hit *OTUD5* has not been previously implicated in the regulation of pluripotency. In both NE and DE differentiation contexts, *OTUD5* KOs exhibited reduced levels of OCT4, SOX2, and NANOG expression (**Fig. 6a-d**). However, under hESC maintenance conditions, *OTUD5* KOs maintained normal levels of these pluripotency markers (**Supplementary data Fig. S9a-c**). This suggests that the lower levels of pluripotency markers observed during differentiation do not stem from lower levels of these factors in hESCs prior to differentiation, but rather arise specifically during the differentiation process. These findings, consistent with the NE and DE screen results, highlight the role of OTUD5 in safeguarding pluripotency upon exposure to differentiation cues, thus ensuring the ordered dissolution of the pluripotent state. However, this loss of pluripotency phenotype doesn’t coincide with an acquisition of differentiation features. While we observed a statistically significant but small increase in the percentage of SOX17+ endoderm cells in *OTUD5* KOs during DE differentiation, there was a significant decrease in the percentage of PAX6+ cells under NE differentiation (**Supplementary data Fig. S9d-f**). Therefore, OTUD5 acts not as a general mediator of differentiation competence, but rather specifically regulates in the dissolution of pluripotency, emphasizing the need to interrogate the regulation of the dissolution of pluripotency separately from that of differentiation.

**Figure 6.**
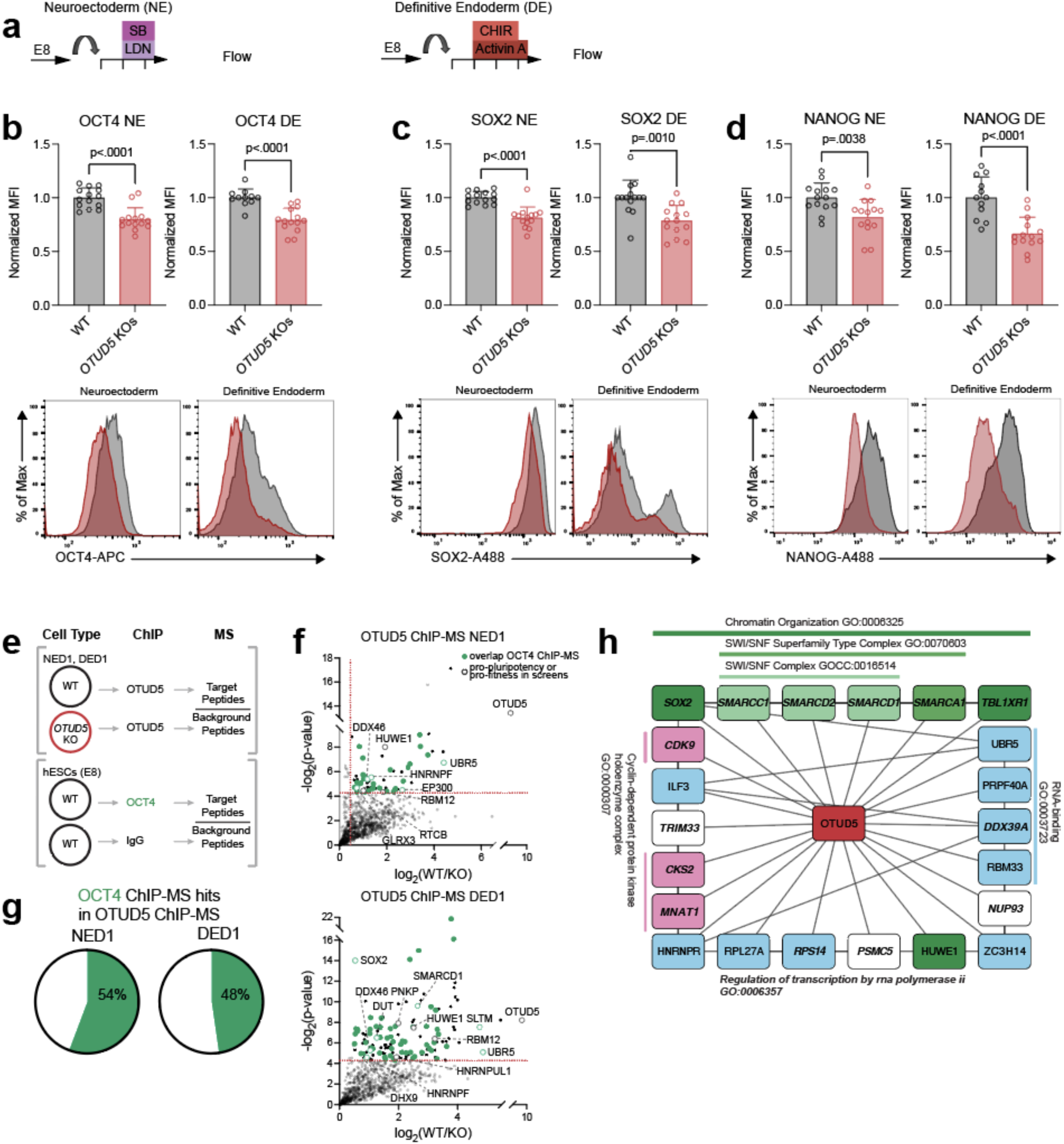
*OTUD5* KOs identify role for OTUD5 in the regulation pluripotency. (a) Schematic of NE and DE differentiations for flow cytometry. (b) Flow cytometry quantification and representative histograms OCT4, (c) SOX2, and (d) NANOG in WT and *OCT4* KO hESCs in NE and DE differentiations. HUES8 iCas9 WT2, WT3, and HUES8iCas9 *OTUD5^-/-^* KO1, and KO2, lines analyzed in each experiment. *n=* 7 independent experiments. Error bars indicate s.d. Statistical analysis was performed by unpaired two-tailed Student’s t-test. P-values are indicated. (e) Schematic of OTUD5 and OCT4 ChIP-MS. (f) Plots showing proteins identified in OTUD5 ChIP-MS in NE Day 1 and DE Day 1. Dotted lines indicate the cut-offs (log2(Fold Change OTUD5/IgG)) > 0.5, –log2 (P) > 4.32) for significantly enriched proteins. Proteins that are also identified by OCT4 ChIP-MS in hESCs are indicated with green circles. Pro-pluripotency and pro-fitness screen hits labelled. OTUD5 ChIP-MS *n =* 3 independent experiments using HUES8 iCas9 WT2, and HUES8iCas9 *OTUD5^-/-^* KO2. OCT4 ChIP-MS *n* = 2 independent experiments using HUES8iCas9 parental hESCs. (g) Proportion of proteins identified by OCT4 ChIP-MS in OTUD5 ChIP-MS hits. (h) Cytoscape network visualization displays the interactions between 22 hits identified by OTUD5 ChIP-MS. To select these hits, we merged the top 15 hits by p-value in NE and DE OTUD5 ChIP-MS, and only included those with interaction scores >0.4 with at least one other hit based on STRING analysis ^37^. Edges and GO terms determined were determined by STRING analysis. Edges visualizing the connection with OTUD5 from the ChIP-MS added.

*OTUD5*, identified in module 8 along with a large number of chromatin regulators, is located in the nucleus ^66–69^, suggesting a role in chromatin regulation. To investigate this further, we performed chromatin immunoprecipitation mass spectrometry (ChIP-MS) to identify chromatin factors associated with OTUD5 during early NE and DE differentiation (day 1) using *OTUD5* KO cells as a negative control (**Fig. 6e-f, Supplementary Table 5**). We found that OTUD5 was associated with multiple known regulators of pluripotency including pro-pluripotency and pro-fitness hits identified in our screens. Notably, this included hits identified in the same module as *OTUD5*: SOX2, EP300, and multiple BAF complex members, the common ATPase of which, SMARCA4, was identified in module 8. A large proportion of OTUD5 ChIP-MS hits were also found to be associated with OCT4, a master pluripotency regulator and member of module 8, according to OCT4 ChIP-MS results (**Fig. 6e-g, Supplementary Table 5**). The top OTUD5 ChIP-MS hits were particularly enriched for regulators of active transcription and chromatin organization (**Fig. 6h**), reflecting positive regulators of pluripotent identity and cell fitness found in our screens. Our findings indicate a previously underappreciated role for open chromatin in the stability and maintenance of the pluripotent cell state and suggest that this process involves OTUD5 and the regulation of ubiquitination. Indeed, *OTUD5* KO resulted in the decrease of markers of pluripotency during differentiation (**Figure 6b-d**). OTUD5 has not been previously linked to pluripotency, our findings are consistent with the observation that knockout of *Otud5* in mice is embryonic lethal ^69,70^, and that OTUD5 regulates open chromatin in neuronal development ^69^.

## Discussion

The self-renewal of PSCs is defined by their ability to be propagated indefinitely while maintaining their pluripotent identity ^71–75^. However, hPSC characterization, especially large-scale screening efforts, have largely focused on the cell fitness aspect of self-renewal ^76^. A fundamental question that remains unanswered is whether pluripotent identity is regulated by the same mechanisms as cell fitness. Further, while pluripotency in development is transient and quickly transitions into the three germ layers, much of the research on human pluripotency has been performed under continuously self-renewing PSC maintenance conditions. Our NE and DE screens interrogate the dissolution of pluripotent identity in dynamic transition states reflective of human development. The identification of shared hits between these NE and DE exit of pluripotency screens suggests the existence of common mechanisms regulating the dissolution of pluripotency independent of external differentiation signals and expands on a current model of pluripotency that is dependent on the balance between differentiation programs ^77^. The prevalence of chromatin regulators in these screens suggests a model of pluripotency beyond regulation by the core pluripotency factors OCT4, NANOG, and SOX2, and emphasizes the significance of dynamic chromatin regulation during the dissolution of the pluripotent state. While multiple epigenetic regulators have been shown to play a role in the maintenance of pluripotency ^78^, our screens in the context of dynamic transition states have identified a large number of regulators of open chromatin, including top hits such as EP300, BAF complex members, SAGA histone acetyltransferase members, and the deubiquitinase OTUD5. Furthermore, by comparing the results from the NE and DE exit of pluripotency screens with those from the E8 and E6 fitness screens, we discovered genes that play distinct roles in cell fitness vs. identity. The *OTUD5* and *TADA2B* gene knockout studies support these findings, indicating that studying pluripotency through the lens of cell fitness alone is insufficient. Characterization of the *TADA2B* KOs and other SAGA complex KOs demonstrate that cell fitness is an unreliable indicator of pluripotent identity. Moreover, in specific conditions, such as observed in the *TADA2B* KOs, cell fitness may be enhanced at the expense of pluripotent identity.

Decoupling acquisition and loss of cell state in dynamic systems is particularly challenging. RNAi screens of iPSC reprogramming in a stage-specific manner showed that acquisition of pluripotency and down-regulation of the somatic cell gene network are distinct events, and both must be completed for successful reprogramming ^79,80^. The dynamic nature of our screens for pluripotent cell identity allowed us to investigate the reverse: whether pluripotency dissolution during NE and DE differentiation involves separate forces that either propel the transitions to new cell states or push the cells away from the pluripotent state. Comparison of our DE context pluripotency screen, using *OCT4* as a readout, with previous DE differentiation screens from our lab, using *SOX17* as a readout ^24^ suggested the pull of differentiation signaling is distinct from the push of pluripotency loss. While top hits from the DE differentiation screen were by and large TFs or developmental signaling pathway components, in our DE pluripotency screen, many of the top hits were chromatin modifiers and transcriptional regulators, genes involved in RNA processing, and ubiquitin modifiers. It is remarkable that we discovered different gene regulatory programs by screens conducted on cells cultured in similar differentiation conditions but utilizing different marker genes, *OCT4* for pluripotency and *SOX17* for DE. The incorporation of more comprehensive single-cells readouts such as scRNA-seq ^81–84^ in future studies will likely provide additional insights into the push and pull of cell-state transitions. Our screens provide an unbiased data set which can inform gene selection for further investigation of the push and pull of cell-state transitions and improvements to simple gene regulatory network models of cell-state transitions ^85^. Characterization of *OTUD5* KOs further illustrates the independent regulation of pluripotent identity and the process of differentiation. Beyond development, understanding the forces regulating the simultaneous acquisition and loss of alternative cell states is vital to the study of metaplasia and cell transformation leading to tumorigenesis. This is particularly important given the differing susceptibility of different lineages to transformation, even in response to the same stimuli ^86^.

To effectively unravel pluripotency regulation, it is necessary to compare screens that assess different aspects of pluripotency, allowing for the identification of various regulatory mechanisms. In our work, we clustered genes based on multiple screen results, enabling the identification of related genes that function in the same complex or regulate the same cellular programs within the same modules. One module was enriched for hits that were positive regulators of pluripotent identity and cell fitness, including the core transcriptional regulators of pluripotency: *OCT4, NANOG,* and *SOX2,* and other known regulators of pluripotency. While clustering of screen results has been used to predict genetic relationships from CRISPR essentiality screens in cancer cell lines ^87^, our results demonstrate that such predictive clustering is possible using a small but targeted number of datasets, and may be further enhanced with incorporation of additional screening data from relevant differentiation contexts. This screening strategy and results may also be applied to understanding more mature stem cells in the stem cell niche. For example, hematopoietic stem cells, while highly proliferative, lose regenerative capacity over time as a result of this proliferation ^88^. The ability to decouple regenerative capacity, driven by cell identity, versus proliferation, is vital for cell engineering and advancing regenerative medicine. As such, the gene sets identified in this study have broad applicability and can be used to identify genes of interest in other biological systems. In hPSCs, understanding the regulation of pluripotency is fundamental to downstream differentiation efforts and the study of human development. The resources generated by this study lay the groundwork for large-scale discovery efforts involving hPSC systems. For example, the NIH MorPhiC consortium aims to characterize knockout phenotypes in diverse human cell types for phenotypic analysis, and these resources will be instrumental in achieving this objective.

## Author Contributions

B.P.R. and D.H. conceptualized the study, designed the experiments, interpreted results. B.P.R. performed most of the experiments and data analyses. Q.V.L. assisted with the screens. H.S.C. performed clustering and Reactome analysis. D.L. and D.Y. performed the cell competition assay. S.G., S.S., J.Y., and R.L. performed ChIP-MS. N.V. generated the OCT4-GFP donor plasmid. J.R.D. and H.K. assisted with characterization of KO lines. S.J.K. performed gRNA enrichment analysis. M.A.B. performed MAGeCK analysis. B.P.R. and D.H. wrote the manuscript. All authors provided editorial advice. D.H. supervised the study.

## Acknowledgements

We thank H. Pan, P. Shila, and J. Giacalone for assisting with additional experiments not included in the manuscript; E. Apostolou and L. Studer for insightful discussions; and the MSKCC Gene Editing and Screening Core and the MSKCC Flow Cytometry Core for technical assistance. This study was supported in part by grants to D.H. from NIH (U01HG012051, UM1HG012654) and New York State Stem Cell Science (NYSTEM, C32593GG), MSKCC Cancer Center Support Grant from NIH (P30CA008748), an NIH T32 Training Grant (T32GM008539 to B.P.R. and S.J.K.), a Howard Hughes Medical Institute (HHMI) Medical Research Fellowship (to N.V.), a NYSTEM training grant from the MSKCC Center for Stem Cell Biology (CSCB) (DOH01-TRAIN3-2015-2016-00006 to D.Y.).

## Methods

### Culture of hESCs

Two human hESC lines were used in this study, H1 (NIHhESC-10-0043) and HUES8 (NIHhESC-09-0021), both with an inducible Cas9 insertion^27,89^. These lines were maintained in chemically defined, serum-free Essential 8 (E8) medium conditions (Thermo Fisher Scientific, A1517001) on tissue culture-treated polystyrene plates coated with vitronectin (Thermo Fisher Scientific, A14700) at 37 °C with 5% CO2. For regular maintenance, hESCs were dissociated with 0.5 mM EDTA (KD Medical, RGE-3130) at a 1:10–1:20 split ratio every 3–5 days. 10 µM Rho-associated protein kinase (ROCK) inhibitor Y-27632 (Selleck Chemicals, S1049) was added into the culture medium when passaging or thawing cells unless otherwise noted. Cells were counted using Vi-CELL XR Cell Viability Analyzer (Beckman Coulter). Cells were routinely confirmed to be Mycoplasma-free by the MSKCC Antibody and Bioresource Core Facility and karyotypically normal by the MSKCC Molecular Cytogenetics Core. All experiments were approved by the Tri-SCI Embryonic Stem Cell Research Oversight Committee (ESCRO).

### Generation of H1 OCT4^GFP/+^iCas9 Reporter Line

#### Generation of OCT4-2A-copGFP donor plasmid

The *OCT4-*GFP reporter construct was constructed using modifications to a selection free knock-in strategy previously used in our lab ^26^. The donor vector for this construct was pOCT4-2A-copGFP. To generate this vector, an NheI-2A-copGFP-AscI cassette was PCR amplified using the primers NhPTV-F and Asc_St_copGFP-R from the plasmid pCRIIAgPTV_ppGFP (An AgeI-PTV1-2A-ppGFP fragment was amplified by PCR using AgPTVppGFP-F and ppGFP-R, using pMAX-GFP plasmid (Lonza) as template. The AgeI-P2A-ppGFP insert was purified and Topo-cloned into pCR™II-TOPO^TM^ (Thermo Fisher Scientific) resulting in pCRIIAgPTV_ppGFP). Next, the NheI-2A-copGFP-AscI was ligated into the pCR^TM^II-TOPO^TM^ cloning vector using the TOPO TA Cloning Kit (Thermo Fisher Scientific, 450641) following manufacturers instructions for ligation and transformation. Next, the pCR™II-TOPO^TM^-NheI-2A-copGFP-AscI plasmid and the pOCT4-2A-copGFP plasmid were digested with NheI and AscI and ligated. Sequences for primers used for PCR and sequencing are listed in the **Supplementary Table 6**.

#### Transfection into H1iCas9

Guide RNA (gRNA) and tracrRNA were ordered from IDT (Alt-R® CRISPR-Cas9 crRNA and #1072532). RNA molecules and plasmid were transiently transfected into hESCs using Lipofectamine 3000 (Thermo Fisher Scientific, L3000001) following manufacturer’s instructions. Briefly, gRNA and tracrRNA were added at a 10 nM final concentration, 5µg donor plasmid was added. gRNA/tracrRNA and Lipofectamine/plasmid were diluted separately in Opti-MEM (Thermo Fisher Scientific, 31985070), mixed, incubated for 15 min at room temperature (RT), and added dropwise to 500,000 freshly seeded iCas9 hESCs in one well of a 24-well plate. Cas9 expression was induced with 2 µg/ml doxycycline one day prior to transfection, the day of transfection, and one day after transfection. GFP positive clones were isolated through FACS and subsequent single cell colony picking. The H1 *OCT4^GFP/+^* iCas9 reporter line is a heterozygous line as confirmed through PCR and DNA sequencing. *OCT4*-GFP reporter fidelity was confirmed by flow-cytometry analysis. gRNA sequences are listed in **Supplementary Table 6**.

### Flow Cytometry

Flow cytometry analyses were performed as previously described ^24^. Antibodies used for flow cytometry are listed in **Supplementary Table 7**. Briefly, cells were dissociated and stained with DAPI for live GFP data collection or fixed and stained with LIVE-DEAD Fixable Violet Dead Cell Stain (Invitrogen; L34955) and corresponding antibodies for data collection using BD LSRFortessa or BD LSRII. Annexin V staining was performed using the PE Annexin V Apoptosis Detection Kit I as per manufacturer’s instructions (BD Biosciences, 559763). Flow cytometry analysis and figures were generated in FlowJo v10. Gating strategy is shown in **Extended Data Fig. 1D**.

### Neuroectoderm Differentiation

Neuroectoderm differentiation performed as previously described ^90^ with modifications. hESC cultures were disaggregated using TrypLE (Life Technologies, 12563-029) for 4 minutes, collected in E8 media, spun at 200 x g for 5 minutes, and resuspended in E8 media. 400,000 cells per well of 6-well plate were seeded on vitronectin (Thermo Fisher Scientific, A14700) with 10 µM Rho-associated protein kinase (ROCK) inhibitor Y-27632 (Selleck Chemicals, S1049) in E8 medium (Thermo Fisher Scientific, A1517001). 24 hours after plating, cells were washed with PBS and exposed to Essential 6 (Thermo Fisher Scientific, A1516401) with 10µM SB431542 (Tocris, 161410) and 500nm LDN193189 (Cedarlane Labs, 04-0074-02). Media changed every 24hrs.

### Definitive Endoderm Differentiation

Definitive Endoderm differentiation performed as previously described ^91^ with modifications. hESC cultures were disaggregated using TrypLE (Life Technologies, 12563-029) for 4 minutes, collected in E8 media, spun at 200 x g for 5 minutes, and resuspended in E8 media. 300,000 cells per well of 6-well plate were seeded on vitronectin (Thermo Fisher Scientific, A14700) with 10 µM Rho-associated protein kinase (ROCK) inhibitor Y-27632 (Selleck Chemicals, S1049) in E8 medium (Thermo Fisher Scientific, A1517001). 24 hours after plating, cells were washed with PBS and exposed to S1/S2 medium supplemented with 20 ng/ml Activin A (Bon-Opus Biosciences; C687-1mg) for 3 days, and CHIR99021 (Stemgent, 04-0004-10) for 2 days (first day, 5 µM; second day 0.5µM). S1/S2 medium was composed of MCDB131 medium (Thermo Fisher Scientific, 10372019) supplemented with 1.5g/L sodium bicarbonate (Research Products International, S22060), 1x Glutamax (Thermo Fisher Scientific, 35050061), 10mM glucose (Sigma-Aldrich, G8769), and 0.5% BSA (LAMPIRE, 7500804). Media changed every 24 hrs.

### Infection and Expansion for Genome Wide CRISPR-Cas9 Screens

The human Brunello gRNA library ^30^, consisting of 76,441 guide RNAs (gRNAs) targeting 19,114 genes (four gRNAs per gene), was produced and tested as previously described ^24^. A minimum of 200-fold library coverage is typically recommended for screens based on basic phenotypes such as cell survival and growth, given the relatively complex nature of our multiple reporter and essentiality screens, we target a 600X library coverage at all steps to maximize sensitivity. 7 Days before the start or our screens, 142 million H1 OCT4^GFP/+^iCas9 cells were infected with the lentiviral library at an MOI of 0.4 in 150-mm plates at a density of 1.67 million per plate (>600-fold library coverage after selection with puromycin). 6 μg/ml protamine sulfate was added concurrently with the virus infection to enhance the infection efficiency. Infected cells were treated with 2 μg/ml doxycycline (Thermo Fisher Scientific, BP26535) (beginning 24 hours after plating) and 0.5 μg/ml puromycin (Sigma-Aldrich, P8833) (beginning 48 hours after plating). 7 days post-infection, cells were treated with TrypLE Select (Thermo Fisher Scientific, 12563029), counted and replated for four individual screens. This was considered Day 0 of screening.

### Genome Wide CRISPR-Cas9 Screens for Pluripotency

#### NE screen

160 million post-infection and selection D0 H1 OCT4^GFP/+^ iCas9 cells were replated in 150-mm plates at a density of 8 million per plate (>600-fold library coverage). 24hrs after plating, cells were switched from maintenance E8 medium to NE differentiation medium (described in subsection *Neuroectoderm Differentiation*). After 36hrs of NE differentiation, cells were dissociated using TrypLE Select and sorted using FACSArias (BD Biosciences), according to GFP expression. GFP+ and GFP− cells were collected in to two pellets per condition, with ∼50 million cells (>600-fold library coverage) collected per condition. Pellets were frozen for subsequent DNA extraction.

#### DE screen

90 million post-infection and selection D0 H1 OCT4^GFP/+^ iCas9 cells were replated in 150-mm plates at a density of 6 million per plate (>600-fold library coverage). 24hrs after plating, cells were switched from maintenance E8 medium to DE differentiation medium (described in subsection *Definitive Endoderm Differentiation*). After 60hrs of DE differentiation, cells were dissociated using TrypLE Select and sorted using FACSArias (BD Biosciences), according to GFP expression. Sorted GFP+ and GFP− cells were collected in to two pellets per condition, with ∼30 million cells (392-fold library coverage) collected per condition. Pellets were frozen for subsequent DNA extraction.

### Genome Wide CRISPR-Cas9 Screens for Cell Fitness

#### E8 screen

Post-infection and selection D0 H1 OCT4^GFP/+^ iCas9 cells were collected as Day 0 samples, with ∼144 million cells (>600-fold library coverage) in two pellets. Pellets were frozen for subsequent DNA extraction. These Day 0 samples were used as the initial timepoint for both E8 and E6 screens. For later timepoint samples, 66.5 million post-infection and selection D0 H1 OCT4^GFP/+^ iCas9 cells were replated in 150-mm plates at a density of 3.5 million per plate (>600-fold library coverage). Cells were expanded and split again at the same cell number and density with TryPLE Select on Day 4 and Day 7 of expansion. On Day 10 of expansion ∼150 million cells (>600-fold library coverage) were collected in two pellets. Pellets were frozen for subsequent DNA extraction.

#### E6 screen

Day 0 samples collected as described above in “*E8 screen*”. For later timepoint samples, 66.5 million post-infection and selection D0 H1 OCT4^GFP/+^ iCas9 cells were replated in 150-mm plates at a density of 3.5 million per plate (>600-fold library coverage). 24 hrs after plating (Day 1) medium was changed to Essential 6 (E6) medium conditions (Thermo Fisher Scientific, A1516401) and cultured in E6 medium for 120hrs, when medium was changed back to E8 (Day 6). Cells were split again at the same cell number and density with TryPLE Select on Day 7 of expansion. On Day 10 of expansion ∼121 million cells (>600-fold library coverage) were collected in two pellets. Pellets were frozen for subsequent DNA extraction.

### gRNA Sequencing

gRNA enrichment sequencing was performed by MSKCC Gene Editing & Screening Core Facility as previously described ^24^ . Briefly, genomic DNA from cell pellets was extracted using the QIAGEN Blood & Cell Culture DNA Maxi Kit (QIAGEN, 13362) and quantified by Qubit (Thermo-Scientific) following the manufacturer’s guidelines. Two-step PCR was performed to amplify gRNA sequences for HiSeq. The first PCR used primer sequences to amplify lentiGuide-puro using ∼510 μg of gDNA (>1000-fold library coverage) per pellet. This PCR was performed using multiple separate 100 μL reactions each with 10 μg gDNA for 18 cycles, with pooling of the resulting amplicons by sample. For the second PCR, 5 μL of product from the first PCR was used in a 100 μL reaction for 24 cycles, with primers to attach Illumina adapters for barcoding. Primers from ^24^. Gel-purified amplicons were quantified by Qubit and Bioanalyzer (Agilent) and sequenced on the Illumina HiSeq 2500 platform. Raw FASTQ files were demultiplexed and further processed to only contain unique gRNA sequences, and the processed reads were aligned to gRNA library sequences using the FASTX-Toolkit (http://hannonlab.cshl.edu/fastx_toolkit/). For each sample, all available reads were combined from different sequencing runs. Read count normalization was performed to median number of reads per sample as part of MAGeCK analysis ^31^.

### Pluripotency Screens Data Analysis

Genes were ranked by gRNA read count using the MAGeCK (model-based analysis of genome-wide CRISPR-Cas9 knockout) robust ranking aggregation (RRA) algorithm ^31^ using MAGeCK 0.5.9.4 default RRA parameters. In each screen, pro-pluripotency hits were defined as genes with 150 lowest ranked RRA scores (OCT4-GFP^lo^ enrichment), anti-per pluripotency hits were defined as genes with 150 lowest ranked RRA scores (OCT4-GFP^hi^ enrichment). Pluripotency scores calculated per screen per gene = log10(RRA score OCT4-GFP^hi^ enrichment) - log10(RRA score OCT4-GFP^lo^ enrichment). Log_2_(Fold Change OCT4-GFP^hi^ / OCT4-GFP^lo^) (LFC) was calculated per gene using MAGeCK 0.5.9.4 default parameters. Screen results are found in **Supplementary Table 1**. For GSEA top hit sets, genes were ranked by pluripotency score and GSEA performed with GSEA Software Version 4.2.3 ^92,93^ using pre-ranked option. Screening data plotted using ggplot2 R-package ^94^ formatted with Adobe Illustrator.

### Comparison to Previous Differentiation Screens

For comparison to other screens, ectoderm differentiation screen data from Naxerova et al. ^9^ based on EpCAM+/NCAM-vs. EpCAM-/NCAM+ enrichment was used. These screens used 2 CRIPSR libraries covering roughly 2/3rs and 1/3^rd^ of coding genome, published MaGECK analysis data for both ectoderm screen sets (“CRISPR Ectoderm P13 Mageck” and “CRISPR Ectoderm P2 Mageck”) were combined, then all genes ranked by published RRA scores. We defined pro-ectoderm differentiation hits as 150 genes with lowest neg. RRA score and anti-ectoderm differentiation hits as 150 genes with lowest pos. RRA score. Endoderm differentiation screen data from Li et al. ^24^ based on SOX17+ and SOX17-enrichment. We used “Brunello MaGECK” data set to define hits, pro-endoderm differentiation hits defined as 150 genes with the lowest pos. RRA score, and anti-endoderm differentiation hits defined as 150 genes with lowest neg. RRA score.

Comparison hit overlaps visualized with Upset plots ^95^ using the online implementation of UpsetR package ^96^ UpSetR Shiny App (https://gehlenborglab.shinyapps.io/upsetr/) which were then formatted with Adobe Illustrator. Analysis of hit lists performed with STRING database v11.5 ^37^ with enriched terms for “STRING clusters”, “GO Component”, and “COMPARTMENTS” shown. GO term enrichment analysis performed using Metascape 3.5 ^38^ (https://metascape.org/).

### Fitness Screen Analysis

Genes were ranked by gRNA read count using the MAGeCK (model-based analysis of genome-wide CRISPR-Cas9 knockout) robust ranking aggregation (RRA) algorithm ^31^ using MAGeCK 0.5.9.4 default RRA parameters. In each screen, pro-fitness hits were defined as genes with 150 lowest ranked RRA scores (Day 0 enrichment), anti-fitness hits were defined as genes with 150 lowest ranked RRA scores (Day 10 enrichment). Pluripotency scores calculated per screen per gene = log10(RRA score Day 10 enrichment) - log10(RRA score Day 0 enrichment). Log_2_(Fold Change Day 0/ Day 10) (LFC) was calculated per gene using MAGeCK 0.5.9.4 default parameters. Screen results are found in **Supplementary Table 1**.

### Comparisons and Clustering Pluripotency and Fitness Screens

Pearson correlation test, UpSet plots, GSEA and STRING analysis performed as described in *Pluripotency Screen Analysis*. Enrichment analysis on the Reactome Pathway Database was performed on top 150 hits from each screen that could be uniquely identified by the Entrez ID using the Reactome-PA R package ^48^. These Reactome sets were then ranked by Median |LFC| of the genes intersected with the top 150 gene group in the screen. For the hierarchical clustering of pro-pluripotency and pro-fitness screening hits, relative levels LFC were represented as column z-scores, and hierarchical clustering was done using the Pearson correlation chosen as the distance metric and Ward’s algorithm as the linkage method. The top 8 distinguished branches in the dendrogram were defined as modules. Modules were further characterized by STRING database V11.5 ^37^ using k-means clustering (k=5) with default parameters.

### Comparisons to Existing Fitness, Pluripotency, and Essentiality Datasets

For comparison to other hESC fitness screens data from ^11–13^ was used, as re-analyzed in ^12^ by BAGEL analysis ^97^. For all fitness screens, top 150 pro-fitness hits were calculated by highest Bayes Factor for fold change in late time point vs. early timepoint. Data for ^12^ screens were taken from the MEF “T12” set, and the Laminin “T12” set. ^11^ data was from the “day30” set, and ^13^ was from the “T18” data set. For the pluripotency screen in hESC maintenance conditions, top 150 pro-pluripotency hits were calculated by RSA score^98^. Data for ^13^ pluripotency screen was taken from the “OCT4_low_high_RSA set. Common essentiality genes shown are the “CRISPRInferredCommonEssentials” data set from DepMap ^50,51,53^ version 22Q4 ^49^. Precision and recall were calculated using pluripotency and fitness scores from NE, DE, E8, and E6 screens, using the essential (625 genes) and non-essential (350 genes) as defined by by Wang et al ^40^. The essential and non-essential sets were used as true positive and true negative lists for PRC using the PRROC R-package ^99,100^.

### Screen Hit Validation

Hit validation was performed using the lentivirus CRISPR approach to generate knockouts in H1 OCT4^GFP/+^iCas9 cells. gRNAs from the Brunello library are listed in **Supplementary Table 6**. gRNAs were cloned into lentiGuide-puro (Addgene, 52963) following published protocols ^101^. The lentiGuide-puro construct expresses a puromycin resistance gene, allowing for the selection of infected cells through puromycin treatment. 1 μg lentiGuide, 0.1 μg pCMV-VSV-G ^102^ (Addgene, 8454), and 0.4 μg psPAX2 (Addgene, 12260) plasmids were transfected with the JetPRIME reagent (VMR, 89137972) into 293T cells to pack lentiviruses. Viral supernatant was collected, aliquoted, and stored at −80 °C. A MOI of 0.30∼0.36 was used for the infection of the H1 *OCT4^GFP/+^* iCas9 cells with different lentiCRISPR viruses 7 days before plating for validation. 6 μg/ml protamine sulfate was added concurrently with the virus infection to enhance the infection efficiency. Reflecting screen conditions, infected cells were treated with 2 μg/ml doxycycline (Thermo Fisher Scientific, BP26535) (beginning 24 hours after plating) and 0.5 μg/ml puromycin (Sigma-Aldrich, P8833) (beginning 48 hours after plating). 7 days post-infection, cells were treated with TrypLE Select (Thermo Fisher Scientific, 12563029), counted and replated for validation. All replicates were performed starting from viral infection.

#### NE validation

For NE validation post-infection and selection H1 OCT4^GFP/+^ iCas9 cells were replated in 6 well plates at 400,000 cells/well. 24hrs after plating, cells were switched from maintenance E8 medium to NE differentiation medium (described in the NE differentiation subsection). After 36hrs of NE differentiation, cells were dissociated using TrypLE Select and GFP levels were analyzed by flow cytometry. Relative intensity of OCT4-GFP = (Mean Fluorescent Intensity (MFI) per gRNA) / (Experimental Mean (MFI non-targeting controls)). All experimental repeats were performed starting from viral infection.

#### DE validation

Given our observation that DE differentiation efficiency is density dependent, we were concerned that varying growth rates of knockout hESCs might affect DE differentiation and therefore the downregulation of pluripotency, even for genes that were not direct regulators of the dissolution of pluripotency in DE context. DE validation was performed with normalization to an in-well uninfected tdTomato (tdT)+ control, given the sensitivity of DE differentiation to variability in plating density. To generate the tdT+ control, H1 OCT4^GFP/+^iCas9 cells were previously infected with virus containing the tdT containing plasmid pWPXL_Luc2tdT ^103^ which was a gift from Wenjun Guo. tdT+ clones were isolated through FACS and subsequent single cell colony picking. For DE validation post-infection and selection H1 OCT4^GFP/+^ iCas9 cells were co-plated with tdT+ cells at 150,000 cells/well of each for 300,000 cells/well total. 24hrs after plating, on D8, cells were switched from maintenance E8 medium to DE differentiation medium (described in the DE differentiation subsection). After 60hrs of DE differentiation (D8-D10.5), cells were dissociated using TrypLE Select and GFP levels were analyzed by flow cytometry. Relative intensity of OCT4-GFP = (MFI per gRNA/ MFI in-well tdT+ control) / (Experimental Mean (MFI non-targeting controls/ MFI in-well tdT+ controls for non-targeting gRNAs)). All experimental repeats were performed starting from viral infection.

#### E8 validation

For E8 validation post-infection and selection H1 OCT4^GFP/+^ iCas9 cells were replated in 6 well plates at 175,000 cells/well (D0). Cells were expanded and split again at the same cell number and density with TryPLE Select on Day 4 and Day 7 of expansion. On Day 10 of expansion, cells were dissociated using TrypLE Select and GFP levels were analyzed by flow cytometry. Relative intensity of OCT4-GFP = (MFI per gRNA) / (Experimental Mean (MFI non-targeting controls)). All experimental repeats were performed starting from viral infection.

#### E6 validation

For E6 validation post-infection and selection H1 OCT4^GFP/+^ iCas9 cells were replated in 6 well plates at 175,000 cells/well (D0). 24 hrs after plating (Day 1) medium was changed to Essential 6 (E6) medium conditions (Thermo Fisher Scientific, A1516401) and cultured in E6 medium for 120hrs, when medium was changed back to E8 (Day 6). Cells were split again at the same cell number and density with TryPLE Select on Day 7 of expansion. On Day 10 of expansion wells were dissociated using TrypLE Select and cells/well counted the using Vi-CELL XR Cell Viability Analyzer (Beckman Coulter). All experimental repeats were performed starting from viral infection.

### Generation of Clonal Knockout hESCs

Clonal knockouts (KOs) were generated in the HUES8 iCas9 hESC ^89^ as previously described with some modifications ^104^. Sequences for of gRNAs and primers used for PCR and sequencing are listed in **Supplementary Table 6**. GRNAs and tracrRNA were ordered from IDT (Alt-R® CRISPR-Cas9 crRNA and #1072532). RNA molecules were transiently transfected into hESCs using Lipofectamine RNAiMAX (Thermo, 13778100) following manufacturer’s instructions. Briefly, gRNA and tracrRNA were added at a 15 nM final concentration. gRNA/tracrRNA and Lipofectamine RNAiMAX were diluted separately in Opti-MEM (Thermo, 31985070), mixed together, incubated for 15 min at room temperature (RT), and added dropwise to 250,000 freshly seeded iCas9 hESCs in a 24-well plate. Cas9 expression was induced with 2 μg/ml doxycycline one day prior to transfection, the day of transfection, and one day after transfection. Three to four days after transfection, hESCs were dissociated to single cells using TrypLE Select (Thermo Fisher Scientific, 12563029), and 500-1000 cells were plated into one 100-mm tissue culture dish with 10 ml E8 media supplemented with 10 μM ROCK inhibitor Y-27632 (Selleck Chemicals, S1049) for colony formation. After 10 days of expansion, 96 colonies were picked into individual wells of a 96-well plate. gDNA from crude cell lysate was used for PCR genotyping, followed by expansion of KO cell lines. Additional sequencing performed on gDNA extracted by QIAGEN Blood & Cell Culture DNA Maxi Kit (QIAGEN, 13362), followed by PCR and insertion into Zero Blunt TOPO PCR Cloning Plasmid (Thermo Fisher Scientific, 450245) which were transfected and expanded per manufacturer instructions. Plasmid was miniprepped using the Zyppy Plasmid Miniprep Kit (Zymo, D4037) per manufacturer’s instructions and sequenced. Clonal KOs were also confirmed by western blot.

### Western Blots

Cell pellets were snap frozen in liquid nitrogen and lysed in cell lysis buffer (9803, Cell Signaling Technology) with proteinase/phosphatase inhibitors (5872, Cell Signaling Technology) and 1 mM PMSF (ICN19538105, MP Biomedicals). Proteins were precleared by centrifugation at 14,000g for 10 min at 4 °C. Protein concentration was determined by the Bradford Protein Assay (Bio-Rad, 500-0202). Equal amounts of protein were loaded into Bis-Tris 10% gel (Novex, NP0301BOX) and transferred to nitrocellulose membranes (Novex, LC2001). Membranes were blocked with 5% milk (LabScientific, M-0841). Primary antibody was incubated overnight at 4 °C. Membranes were washed with TBST three times for 10 min each and incubated with fluorescent conjugated secondary antibody for 1 h at room temperature. Membranes were washed with TBST three times for 10 min each. Blots were visualized using the Odyssey DLx Imaging System (LICOR) Antibodies used for western are listed in **Supplementary Table 7.**

### Growth Curves

hESCs were disaggregated using TrypLE Select and then mechanically dissociated into single cells using 1000 µl tips. One hundred thousand cells were plated into one well of a 6-well plate on vitronectin in E8 medium with ROCK inhibitor. Cells were subsequently maintained in E8 and harvested after TrypLE treatment every 24 hrs for counting cell numbers. For E6 growth curves, cells were rinsed with PBS and changed to Essential 6 (Thermo Fisher Scientific, A1516401) 24 hrs after plating. For -ROCKi growth curves, cells were plated in E8 without ROCK inhibitor.

### Cell Competition Assay

Individual LARRY barcode constructs were cloned from the LARRY barcode library (Addgene:140024) ^105^ and transfected to 293T cells to generate lentivirus. Next, each *OTUD5* KO, and WT clone was infected with a unique LARRY barcode at MOI ∼0.3. One week after lentiviral infection, the barcoded *OTUD5* KO and WT cells, which expressed GFP, were isolated by FACS. Sorted cells were then expanded for 2-3 passages in E8 medium. To do the cell competition test, an equal number of barcoded OTUD5 KO and WT cells were pooled and seeded in 6-well plates 200k per well. Cells were passaged every 3-4 days by TrypLE dissociation and 200k cells were seeded every time. 1, 2, 3 and 4 weeks after pooling, cells were collected for genomic DNA extraction using the Qiagen DNeasy Blood & Tissue Kit (QIAGEN, 69506). The LARRY barcodes were amplified via PCR using the Q5 High-Fidelity DNA Polymerase Kit (NEB, MO0491L) using 500ng genomic DNA and LARRY-F/R primers, sequences are listed in **Supplementary Table 5**. PCR cleanup was performed using AMPure XP Beads (NEB, E7530). A second round of PCR using this purified PCR product using adapters and indexes as described in ^106^. Samples were pooled and submitted to the MSKCC Integrated Genomics Operation where sample quantity and purity were determined using a Qubit fluorometer. Library efficiencies were confirmed by Bioanalyzer (Agilent) and libraries were sequenced on the Illumina HiSeq 400 Platform in PE150 mode, 2-3 million reads per sample. We used CRISPResso2 (http://crispresso.pinellolab.org/submission) ^107^ to quantify the representation of each barcode and thereby each cell line. As each cell-line was labelled with a single bar-code, cell-line representation was calculated by %individual-bar-code in total bar-code reads.

### RNA Isolation and RNA-Seq

Total RNA was extracted using Quick-RNA MiniPrep kits (ZYMO research; R1055) following the manufacturer’s guidelines. Bulk RNA-Seq was performed by the MSKCC Integrated Genomics Operation as previously described ^28,108^ Alignment was performed as described ^109^. DESeq2 ^110^ was used to analyze gene differential expression by comparing transcriptomes of WT and *TADA2B* KO cells in hESCs. DEGs were identified based on cut-off log2(FC) > 1 and false discovery rate (FDR) < 0.05. Results plotted with Enhnced Volcano R-package. GSEA performed with GSEA_4.0.3 using the pre-ranked option and log2(FC) for pairwise comparisons.

### ChIP-MS and Analysis

OTUD5 ChIP-MS was performed on NE and DE Day 1 cells (differentiation described in the NE and DE differentiation subsections) using HUES8iCas9 WT2 and HUES8iCas9 *OTUD5^-/-^* KO2 hESC lines. OCT4 ChIP-MS was performed using HUES8iCas9 hESCs grown in E8 conditions. ChIP-MS was performed from 15 million cells/experiment as previously described ^28,108^. Antibodies used for immunoprecipitation are listed in **Supplementary Table 7**. Proteins were eluted from the ChIP immunoprecipitation using a buffer containing 5% SDS (Thermo Fisher Scientific, AM9820) , 5 mM DTT (Thermo Fisher Scientific, FERR0861) and 50 mM ammonium bicarbonate (pH = 8), and left on the bench for about 1 hour for disulfide bond reduction. Samples were then alkylated with 20 mM iodoacetamide (VWR, IC10035105) in the dark for 30 minutes. Afterward, phosphoric acid (Thermo Fisher Scientific, A2421) was added to the sample at a final concentration of 1.2%. Samples were diluted in six volumes of binding buffer (90% methanol and 10 mM ammonium bicarbonate, pH 8.0). After gentle mixing, the protein solution was loaded to an S-trap filter (Protifi, C02-micro-80) and spun at 500 g for 30 sec. The sample was washed twice with binding buffer. Finally, 1 µg of sequencing grade trypsin (Promega, V5111), diluted in 50 mM ammonium bicarbonate, was added into the S-trap filter and samples were digested at 37°C for 18 h. Peptides were eluted in three steps: (i) 40 µl of 50 mM ammonium bicarbonate, (ii) 40 µl of 0.1% TFA and (iii) 40 µl of 60% acetonitrile and 0.1% TFA. The peptide solution was pooled, spun at 1,000 g for 30 sec and dried in a vacuum centrifuge. Prior to mass spectrometry analysis, samples were desalted using a 96-well plate filter (Orochem) packed with 1 mg of Oasis HLB C-18 resin (Waters). Briefly, the samples were resuspended in 100 µl of 0.1% TFA and loaded onto the HLB resin, which was previously equilibrated using 100 µl of the same buffer. After washing with 100 µl of 0.1% TFA, the samples were eluted with a buffer containing 70 µl of 60% acetonitrile and 0.1% TFA and then dried in a vacuum centrifuge.

Samples were then resuspended in 10 µl of 0.1% TFA and loaded onto a Dionex RSLC Ultimate 300 (Thermo Scientific), coupled online with an Orbitrap Fusion Lumos (Thermo Scientific). Chromatographic separation was performed with a two-column system, consisting of a C-18 trap cartridge (300 µm ID, 5 mm length) and a picofrit analytical column (75 µm ID, 25 cm length) packed in-house with reversed-phase Repro-Sil Pur C18-AQ 3 µm resin. To analyze the proteome, peptides were separated using a 90 min gradient from 4-30% buffer B (buffer A: 0.1% formic acid, buffer B: 80% acetonitrile + 0.1% formic acid) at a flow rate of 300 nl/min. The mass spectrometer was set to acquire spectra in a data-dependent acquisition (DDA) mode. Briefly, the full MS scan was set to 300-1200 m/z in the orbitrap with a resolution of 120,000 (at 200 m/z) and an AGC target of 5x10e5. MS/MS was performed in the ion trap using the top speed mode (2 secs), an AGC target of 1x10e4 and an HCD collision energy of 35. Proteome raw files were searched using Proteome Discoverer software (v2.4, Thermo Scientific) using SEQUEST search engine and the SwissProt human database (updated June 2021). Each analysis was performed with three biological replicates. The search for total proteome included variable modification of N-terminal acetylation, and fixed modification of carbamidomethyl cysteine. Trypsin was specified as the digestive enzyme with up to 2 missed cleavages allowed. Mass tolerance was set to 10 pm for precursor ions and 0.2 Da for product ions. Peptide and protein false discovery rate was set to 1%. Following the search, data was processed as previously described ^111^. Briefly, proteins were log2 transformed, normalized by the average value of each sample and missing values were imputed using a normal distribution 2 standard deviations lower than the mean. Statistical regulation was assessed using heteroscedastic T-test (if p-value < 0.05). Data distribution was assumed to be normal but this was not formally tested. Interaction score and GO analysis for select hits determined by STRING online database ^37^ with visualization of network by Cytoscape ^112^.

### Statistical analysis

All datapoints refer to biological repeats. No statistical method was used to predetermine sample sizes. The investigators were not blinded to allocation during experiments and outcome assessment. No data were excluded from analyses. The number of biological replicates are reported in the legend of each figure. Flow cytometry analysis and growth curves were derived from at least three independent experiments per cell line unless specified in the legends. For ChIP-MS of OCT4 quantification and statistics were derived from two independent experiments. ChIP-MS of OTUD5 quantification and statistics were derived from three independent experiments. CRISPR-Cas9 screening were performed once. All the statistical analyses methods are indicated in the figure legends and methods. Quantification of flow cytometry and growth curve data are shown as the mean ± s.d. Student’s *t*-test was used for comparison between two groups. Statistical significance is indicated in each figure.

### Data Availability

Screening and Mass-Spectrometry in processed and unprocessed form is available in the supplementary tables. Codes are available from the corresponding author upon request. All other data supporting the findings of this study are available from the corresponding authors upon reasonable request.

### Code Availability

Standard scripts and pipelines are described in the methods section of the manuscript. All code supporting the findings of this study are available from the corresponding authors upon reasonable request.

## Supplementary Tables

**Supplementary Table 1.** MAGeCK analysis of NE, DE, E8, and E6 screens. Sheets 1-4 results from MAGeCK ^31^ analysis of NE, DE, E8, and E6 screens. Data were analyzed with MAGeCK 0.5.9.4 default RRA parameters. Sheet 5 top gene-sets as determined by RRA score. Sheets 6-9 NE, DE, E8, and E6 screen gRNA raw counts.

**Supplementary Table 2.** Summary of validation results by hit type.

**Supplementary Table 3.** Previous pro-fitness, pluripotency, essentiality, and differentiation data sets. Sheet 1 gene sets used for GSEA comparison with previous hESC fitness screens. Gene sets from previous screens by Yilmaz et al. ^11^, Mair et al. ^12^, and Ihry et al. ^13^, as re-analyzed by Mair et al. using BAGEL analysis ^97^. For all fitness screens, top 150 pro-fitness hits were calculated by highest Bayes Factor for fold change in late time point vs. early timepoint. Hits are listed based on rank by Bayes Factor. Sheet 2 gene sets from previous screen by Ihry et al. ^13^ using an *OCT4-*GFP reporter during hESC maintenance. Hits are listed based on rank by RSA score. Sheet 3 gene sets from previous Neuroectoderm and Definitive Endoderm screens from Naxerova et al ^9^ and Li et al ^24^. Top 150 pro-differentiation and anti-differentiation were calculated by published RRA score. Hits are listed based on rank by RRA score. Sheet 4 essential and non-essential gene sets used for PRC analysis, determined by analysis from multiple CRISPR screens by Wang et al.^40^ Sheet 5 common essentiality genes shown are the “CRISPRInferredCommonEssentials” data set from DepMap ^50,51,53^ version 22Q4 ^49^. Sheets 6-12 screen results published in Mair et al. ^12^, Ihry et al. ^13^, Naxerova et al ^9^, and Li et al ^24^ , added columns indicate ranking used to determine gene sets.

**Supplementary Table 4.** Pro-pluripotency and pro-fitness hits by module. Top 150 pro-pluripotency and pro-fitness hits from NE, DE, E8, and E6 screens by module, z-scores and

LFC per gene. Z-scores = column scaled LFC. LFC = Log_2_(Fold Change OCT4-GFP^hi^ / OCT4-GFP^lo^) for NE and DE screens or Log_2_(Fold Change Day 0/ Day 10) for E8 and E6 screens.

**Supplementary Table 5.** Results from OTUD5 and OCT4 ChIP-MS analysis. Sheets 1-2 results from OTUD5 ChIP-MS in NE Day 1 (NED1) and DE Day 1 (DED1). *OTUD5* KO cells stained with OTUD5 antibody was used as a negative control. Sheet 3 results from OCT4 ChIP-MS in hESCs (E8). WT cells stained with IgG was used as a negative control. Sheets 4-6, raw data from OTUD5 and OCT4 ChIP-MS experiments.

**Supplementary Table 6.** List of gRNA sequences and primers. Guide RNAs were used for targeting of the *OCT4* locus for creation of the OCT4-GFP reporter line; lentiviral KOs during validation of screening hits; and clonal KO of *OTUD5* or *TADA2B*. Primers used for cloning OCT4-GFP reporter, PCR and sequencing of cell lines, and CRISPR screen Hi-Seq.

**Supplementary Table 7.** List of antibodies

---

**Supplementary Data Figure S1.**
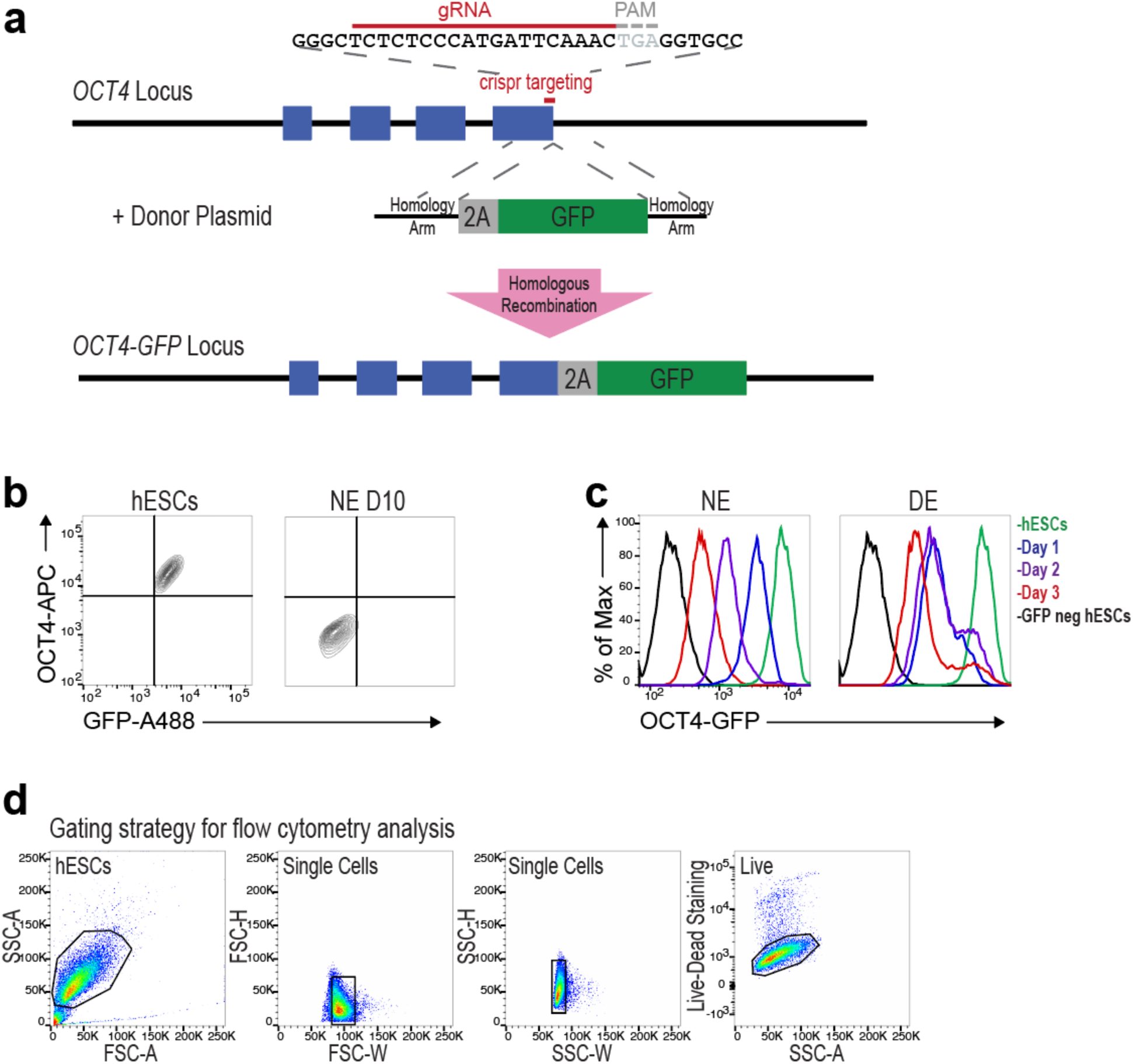
Generation of the *OCT4*-GFP reporter line. (a) Knockin strategy for generating H1 *OCT4^GFP/+^*iCas9 hESCs. (b) Representative flow cytometry contour plots of *OCT4^GFP/+^* hESCs in the maintenance condition (hESC) and on day 10 of neuroectoderm differentiation (NE D10) co-stained for OCT4 and GFP. Representative images are from 2 independent experiments. (c) Representative flow cytometry histograms of time-course of NE and definitive endoderm (DE) differentiations from *OCT4^GFP/+^* hESCs. *OCT4 ^GFP/+^* hESCs were used as positive controls, and H1 iCas9 hESCs (GFP neg hESCs) were used as negative controls. Representative histograms are from 3 independent experiments. (d) Flow cytometry gating strategy. The FSC-A/SSC-A gate identifies cells based on the size and granularity. The FSC-H/FSC-W and SSC-H/SSC-W gates identify single cells. Live-dead staining distinguishes live cells from dead cells.

**Supplementary Data Figure S2.**
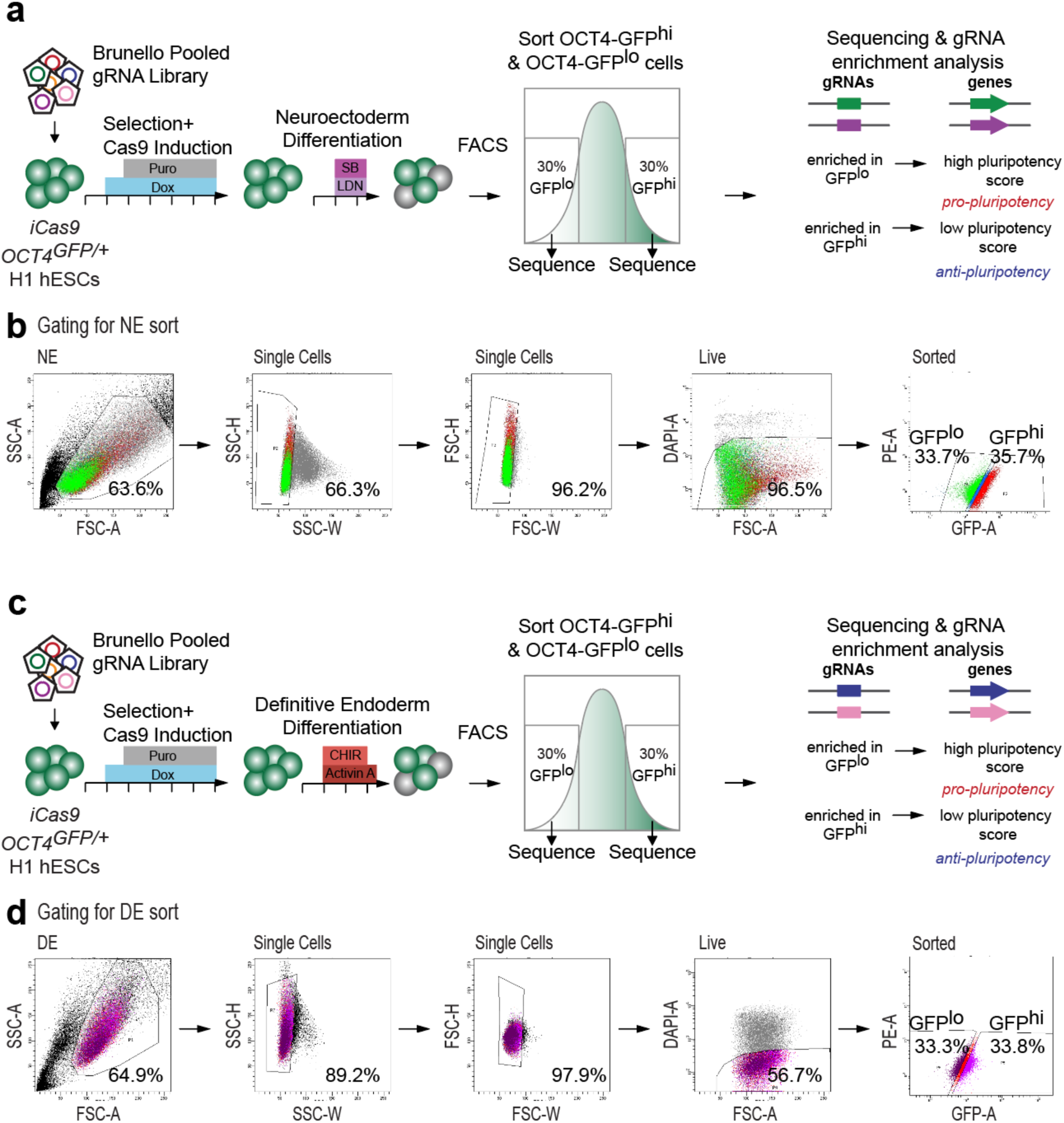
Genome-scale screens to identify regulators of the dissolution of pluripotency in neuroectoderm (NE) and definitive endoderm (DE) contexts. (a) NE CRISPR screen schematic. Each line segment on the horizontal arrows indicates 1 day of media and chemical treatment. (b) Gating used for flow-sorting GFP^hi^ and GFP^lo^ in NE screen (c) DE CRISPR screen schematic and (d) gating.

**Supplementary Data Figure S3.**
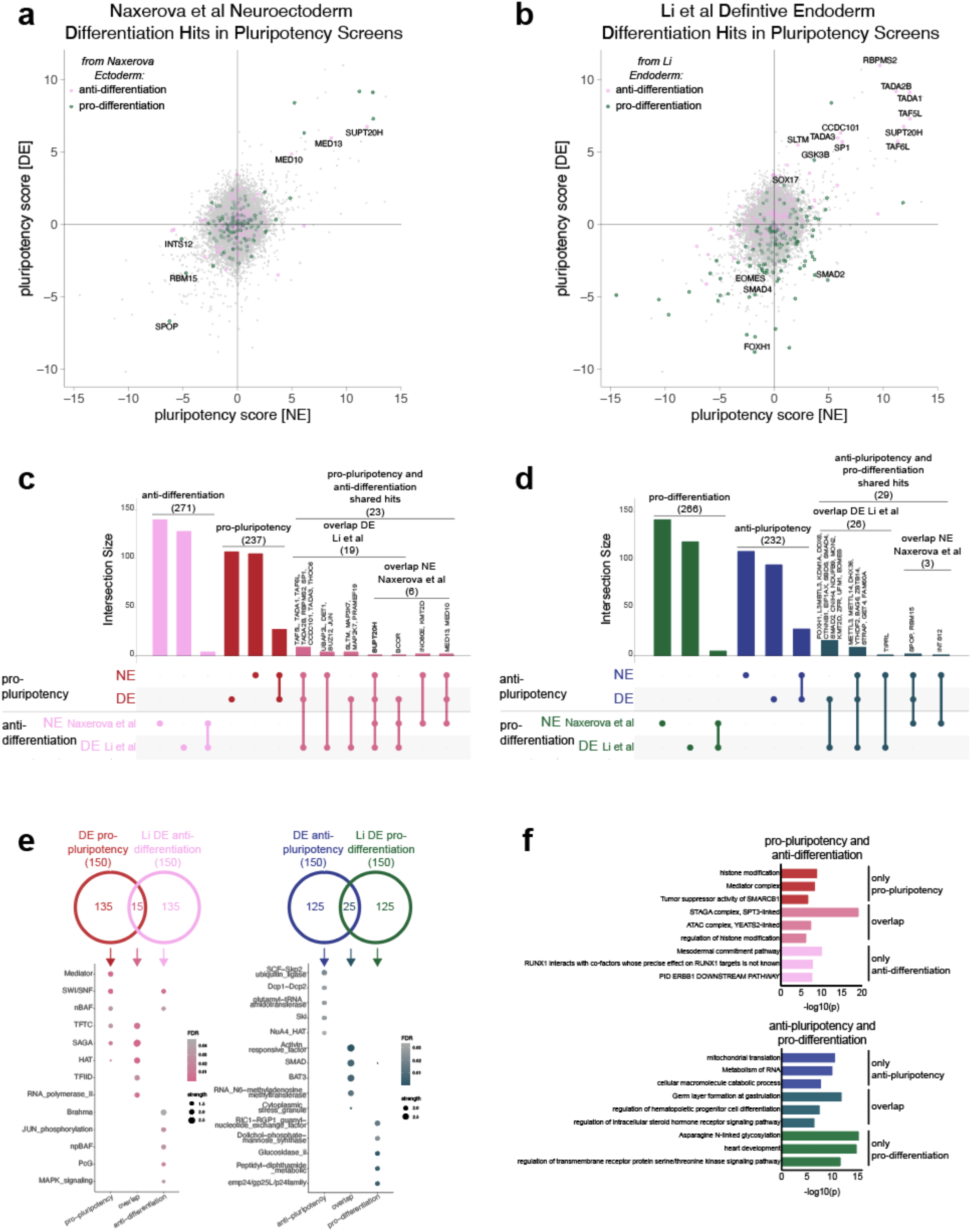
Exit of pluripotency versus differentiation competence screens reveal distinct regulatory networks. (a) Top 150 pro-ectoderm and top 150 anti-ectoderm hits from Naxerova et al. ^9^ (ranked by RRA scores for enrichment in EpCAM+/NCAM- and EpCAM-/NCAM+ populations respectively) in NE and DE screens for pluripotency. (b) Top 150 pro-endoderm and top 150 anti-endoderm hits from Li et al. ^24^ (ranked by RRA scores for enrichment in SOX17- and SOX17+ populations respectively) in NE and DE screens for pluripotency. (c) Overlap of NE and DE pro-pluripotency hits, anti-ectoderm ^9^ hits, and anti-endoderm ^24^ hits as shown by Upset plot ^95^. (d) Overlap of NE and DE anti-pluripotency hits, pro-ectoderm ^9^ hits, and pro-endoderm ^24^ hits as shown by Upset plot ^95^. (e) STRING database analysis overlapping and non-overlapping hits from DE pluripotency screen and Li et al endoderm screen, strength = log10(observed/expected) for enrichment of gene types within a set. (f) GO term enrichment analysis as performed by Metascape ^38^ of overlapping and non-overlapping hits from DE pluripotency screen and endoderm screen.

**Supplementary Data Figure S4.**
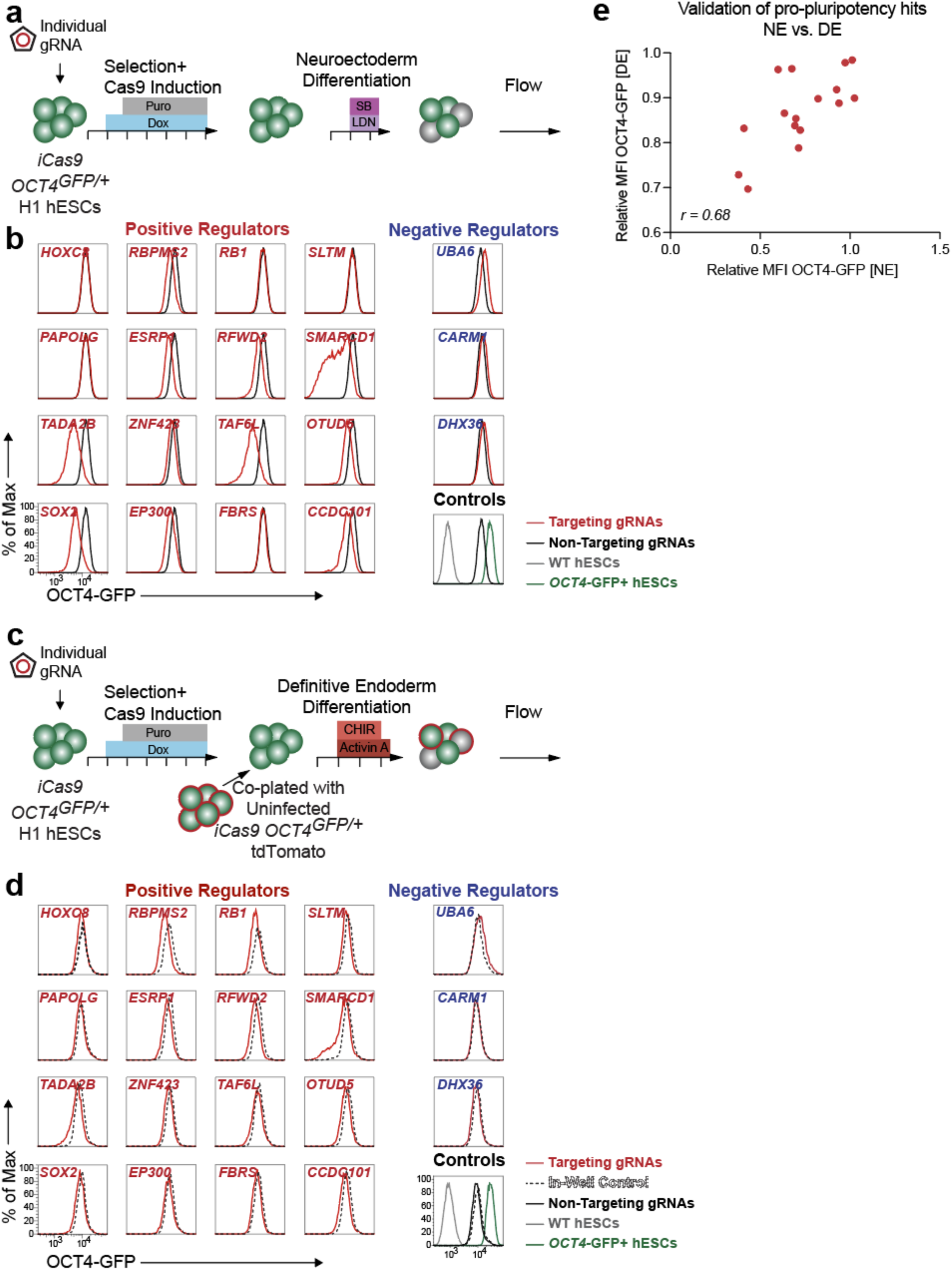
Validation of NE and DE exit of pluripotency screen hits. (a) Schematic of hit validation in NE context using lentiCRISPR infection of H1 iCas9 *OCT4^GFP/+^* hESCs. (b) Flow cytometry histograms show fluorescence OCT4-GFP following gRNA targeting in NE context. Representative images from one of six independent experiments. (c) Schematic of hit validation in DE context using lentiCRISPR infection of H1 iCas9 *OCT4^GFP/+^* hESCs, including in-well uninfected H1 iCas9 *OCT4^GFP/+^* tdTomato (tdT)+ control. (d) Flow cytometry histograms show fluorescence OCT4-GFP following gRNA targeting in DE context. Representative images from one of five independent experiments. (e) Comparison NE context validation to DE context validation. Relative MFI OCT4-GFP [NE] = (MFI per gRNA) / (MFI non-targeting controls). Mean *n*=3-6. Relative MFI OCT4-GFP [DE] = (MFI per gRNA/ MFI in-well tdT uninfected control) / (MFI non-targeting controls/ MFI in-well tdT uninfected control). Mean *n*=5.

**Supplementary Data Figure S5.**
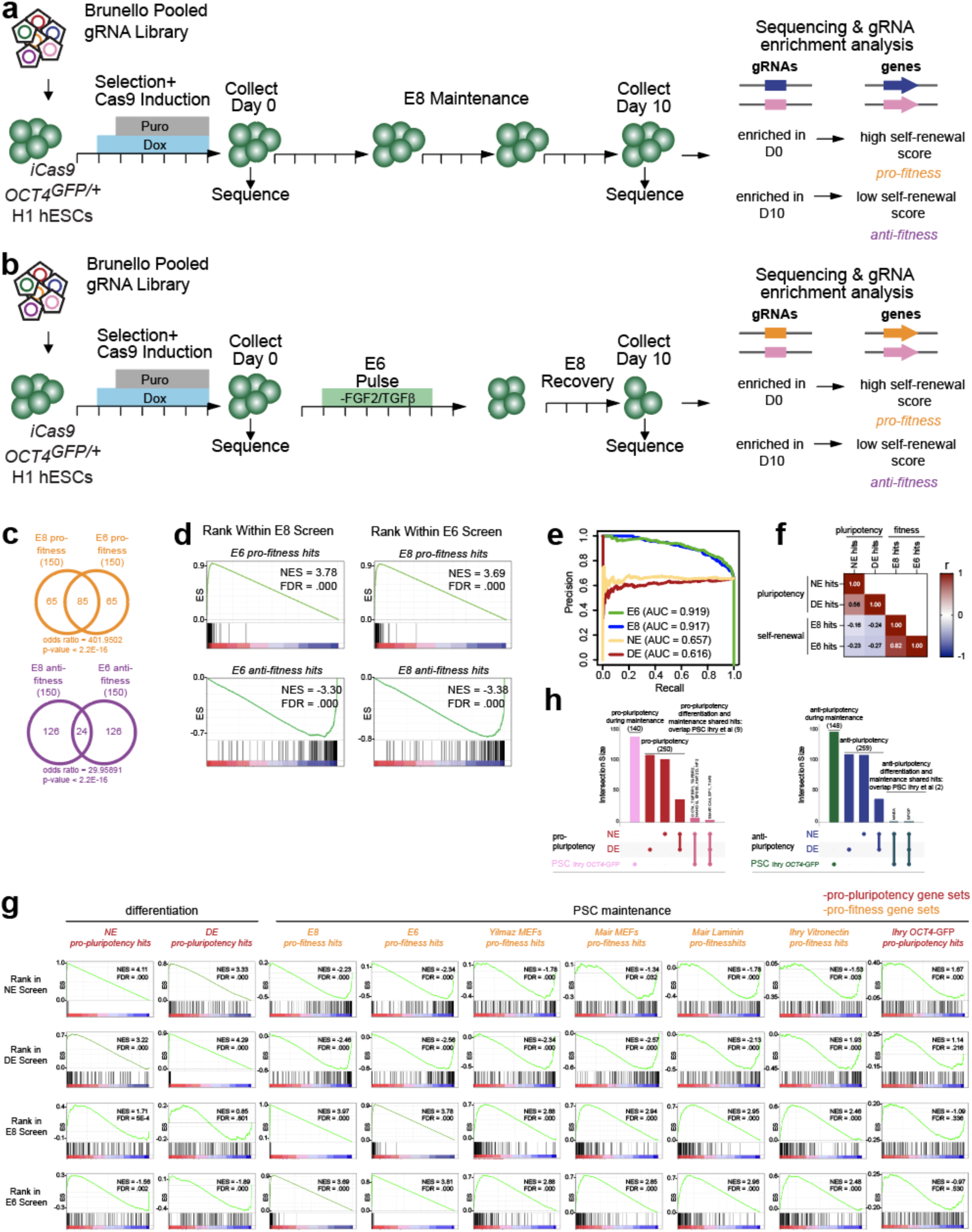
E8 and E6 fitness screens reveal common hits consistent with previous screens for hPSC fitness. (a) E8 CRISPR screen schematic. Each line segment on the horizontal arrows indicates 1 day of media and chemical treatment. (b) E6 CRISPR screen schematic. (c) Overlap of top 150 pro-fitness genes identified in E8 vs. E6 context screens, and top 150 anti-fitness genes identified in E8 vs. E6 context screens, Fisher’s exact test. (d) GSEA for enrichment of pro- and anti-fitness top hits in screen results ranked by fitness score, E8 vs. E6 screens. (e) Precision-recall curves of NE, DE, E8, and E6 screens based on positive RRA scores of essential and non-essential gene sets as defined by Wang et al.^40^ (f) Pearson correlation tests of hits by pluripotency or fitness score. (g) details of GSEA for enrichment of pro-pluripotency and pro-fitness gene sets from NE, DE, E8, E6 screens, previous screens by Yilmaz et al. ^11^, Mair et al. ^12^, and Ihry et al. ^13^, as re-analyzed by Mair et al., and previous pluripotency screen as published in Ihry et al. ^13^. Gene sets from previous fitness screens are top 150 genes per screen as ranked by Bayes factor score, calculated by BAGEL analysis ^97^. Gene sets from previous pluripotency screen are top 150 genes per screen as ranked by RSA score ^98^. Pluripotency screens labelled in red text, fitness screens labelled in orange text. (h) Upset plots ^95^ of overlap NE, DE pro-pluripotency hits and Ihry et al. ^13^ pro-pluripotency hits in maintenance conditions, and overlap of NE, DE anti-pluripotency hits and Ihry et al. ^13^ anti-pluripotency hits in maintenance conditions.

**Supplementary Data Figure S6.**
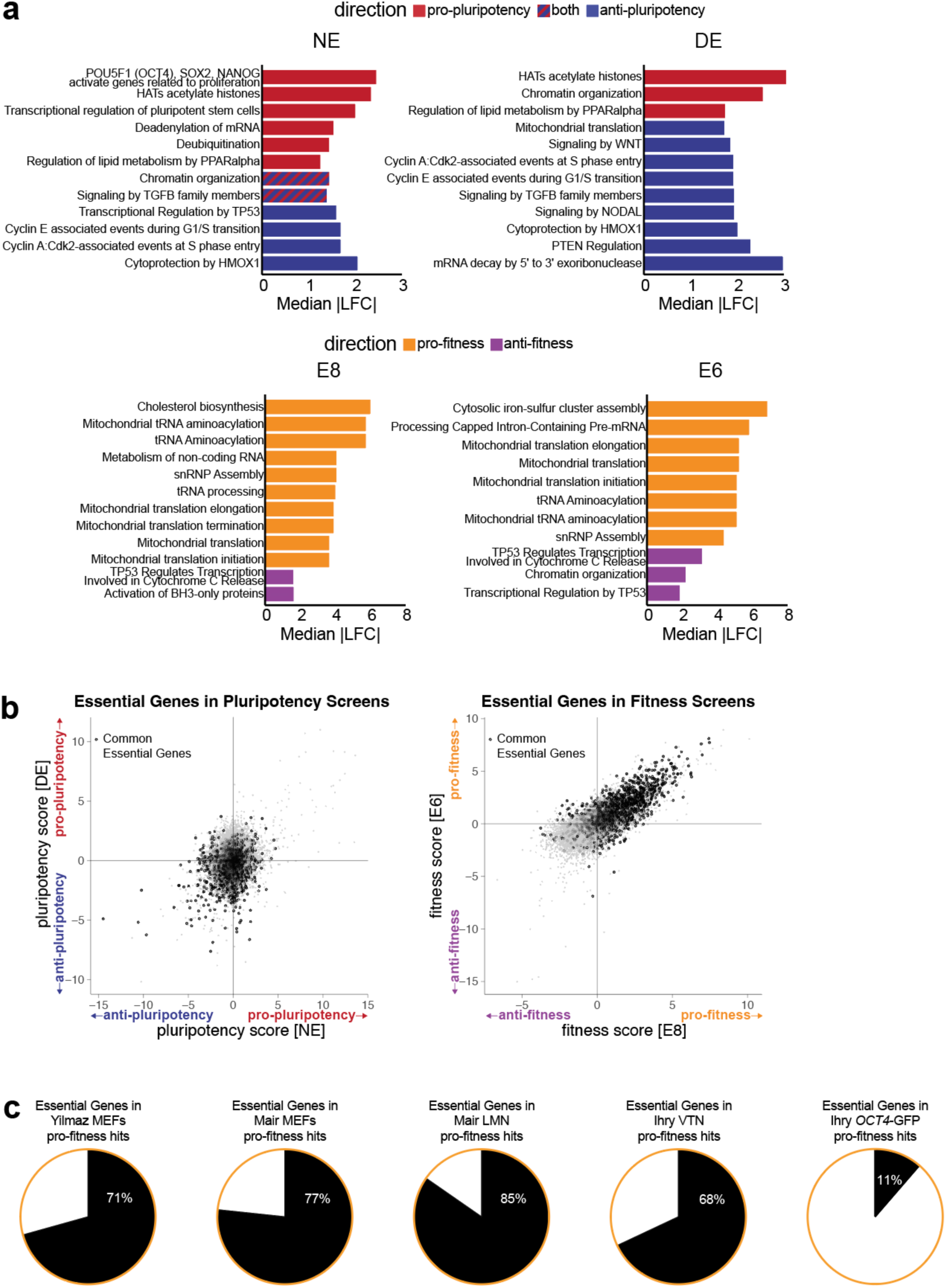
Distinct regulation of pluripotency and hPSC fitness. (a) Top reactome pathway analysis gene sets which score in the pro direction, anti direction, or both. Gene sets identified in top hits of pro-pluripotency, anti-pluripotency, pro-fitness, and anti-fitness NE, DE, E8, and E6 screen hits. Median |Log_2_(Fold Change of OCT4-GFP^lo^/OCT4-GFP^hi^)| or |Log_2_(Fold Change of Day 0/Day 10)| (LFC) from screen calculated for hits identified in reactome set. Highest absolute median |LFC| sets shown. For gene sets identified in both pro and anti directions, mean of median |LFC| calculated for both sets shown. (b) Common essentiality genes by pluripotency score in NE and DE screens, and fitness score in E8 and E6 screens. (c) Proportion of common essentiality genes identified in pro-self renewal gene sets from previous screens by Yilmaz et al., Mair et al., and Ihry et al., as re-analyzed by Mair et al, ^11–13^ and previous pluripotency screen as published in Ihry et al. ^13^. Gene sets from previous fitness screens are top 150 genes per screen as ranked by Bayes factor score, calculated by BAGEL analysis ^97^. Gene sets from previous pluripotency screen are top 150 genes per screen as ranked by RSA score ^98^.

**Supplementary Data Figure S7.**
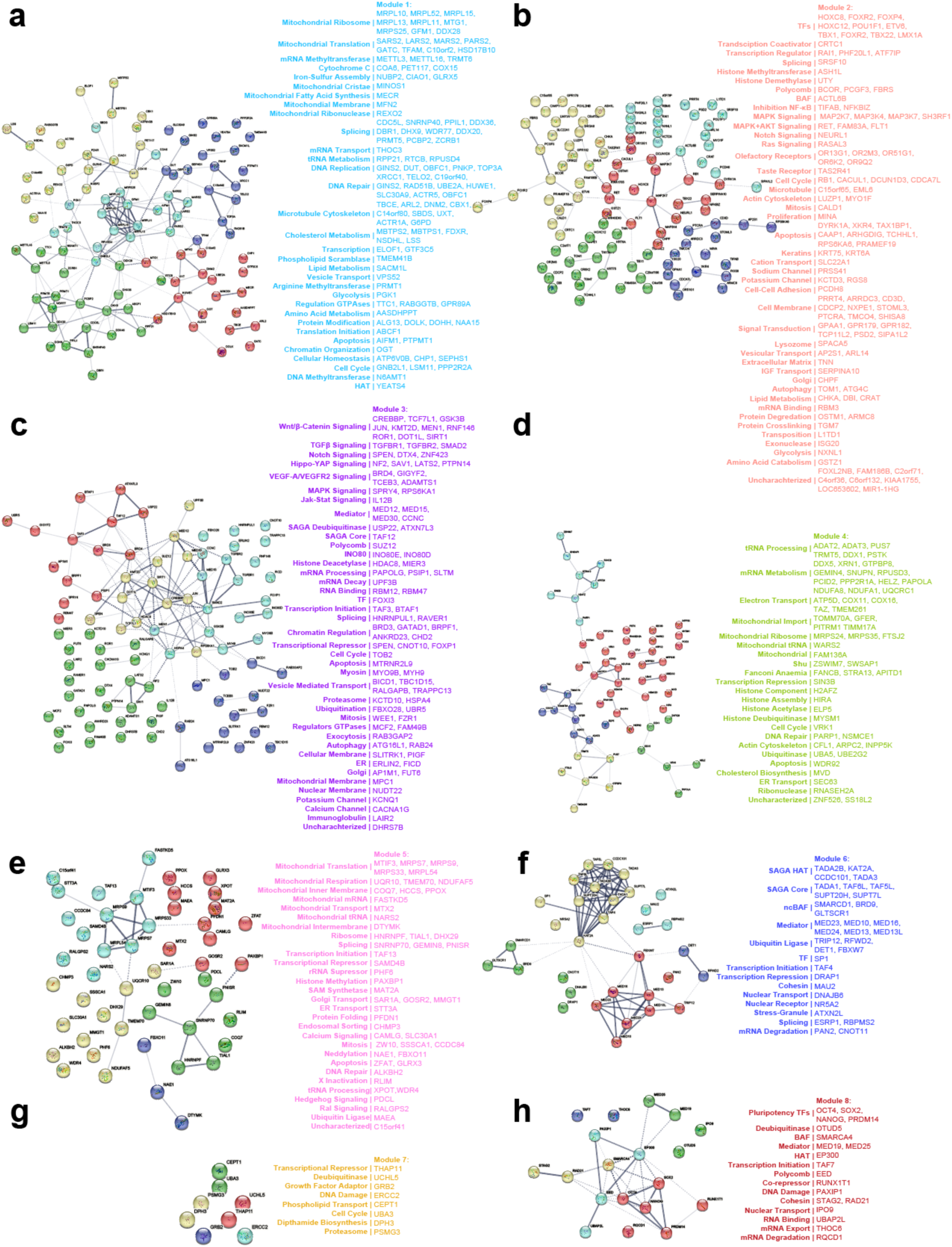
Distinct gene sets identified in individual modules. (a) Interactome indicating known associations between members of cluster gene sets identified through STRING analysis^37^ using default parameters for high-confidence interactions. Each gene-set was further clustered using k-means clustering (k=5), and dotted lines indicate connections between clusters. Annotation of all genes in Module 1, (b) Module 2, (c) Module 3, (d), Module 4, (e) Module 5, (f) Module 6, (g) Module 7, and (h) Module 8.

**Supplementary Data Figure S8.**
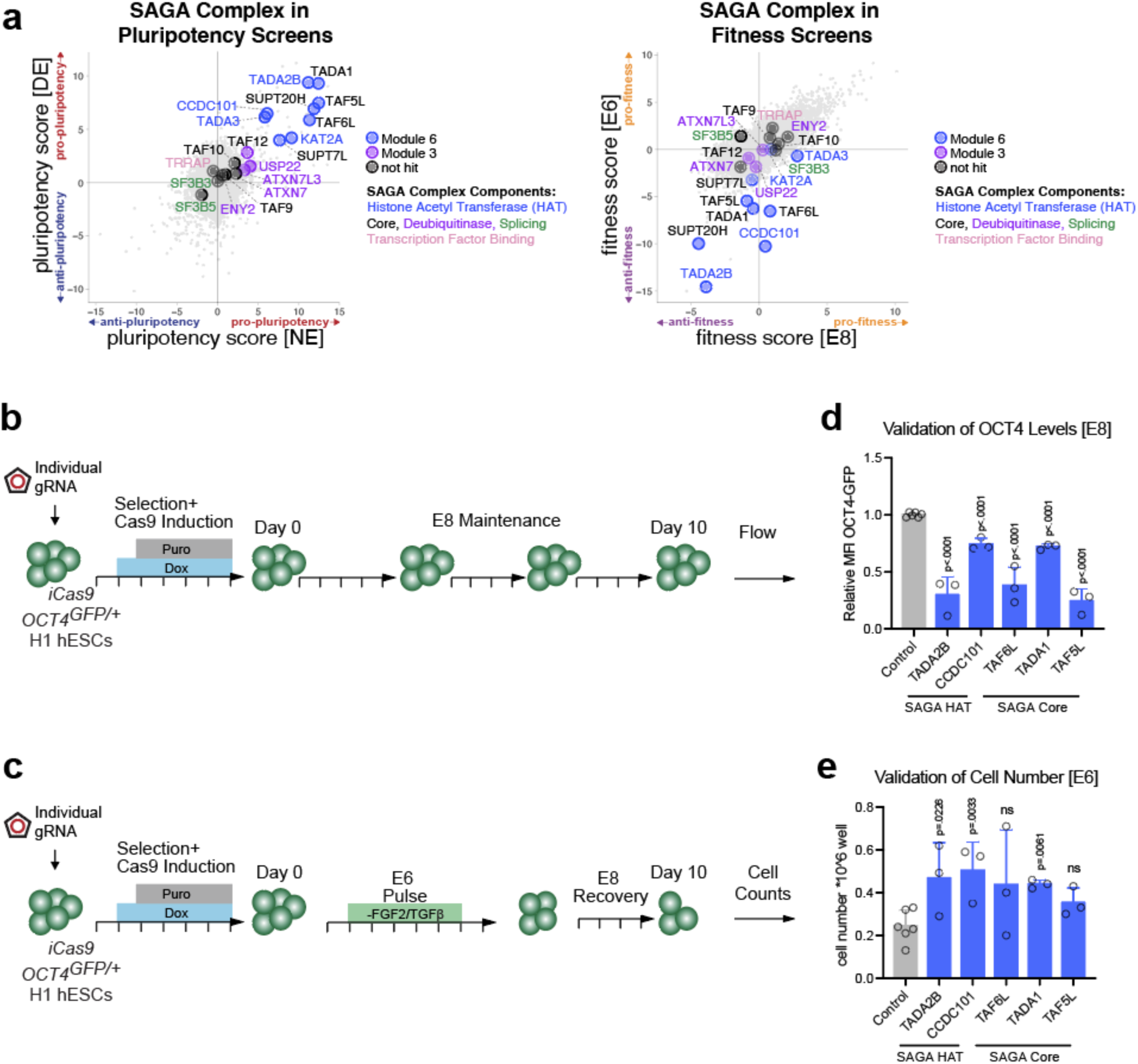
Validation of SAGA complex screen hits reflect *TADA2B* KO pluripotency and fitness phenotypes. (a) SAGA complex components by pluripotency score in NE and DE screens, and fitness score in E8 and E6 screens. (b) Schematic of SAGA complex hit validation of OCT4 levels in E8 maintenance conditions and (c) proliferation in E6 challenge conditions, using lentiCRISPR infection in H1 iCas9 *OCT4^GFP/+^* hESCs. (d) Bar graphs show relative MFI of OCT4-GFP following gRNA targeting in E8 context. Relative MFI OCT4-GFP = (MFI per gRNA) / (MFI non-targeting controls). (e) Bar graphs show cell counts at D10 in E6 context. *n*=3 independent experiments. 2 non-targeting controls analyzed per experiment. Data represented as mean. Error bars indicate s.d. Statistical analysis was performed by unpaired two-tailed Student’s *t*-test. P-values indicated.

**Supplementary Data Figure S9.**
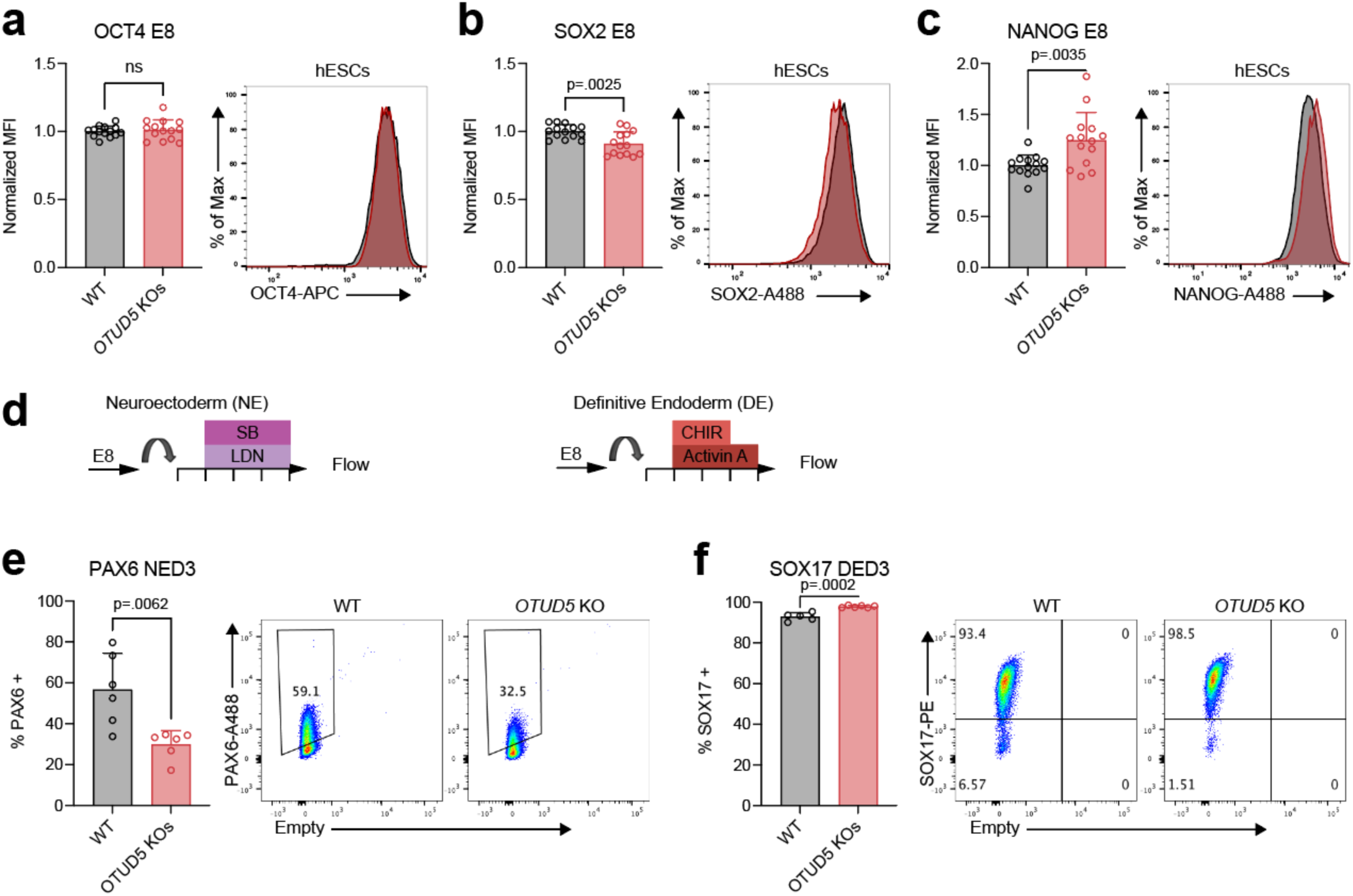
*OTUD5* KOs sensitive to challenges to pluripotency. (a) Flow cytometry quantification and representative histograms OCT4, (b) SOX2, and (c) NANOG in WT and *OCT4* KO hESCs. HUES8iCas9 WT2, WT3, and HUES8iCas9 *OTUD5^-/-^*KO1, and KO2 clonal lines analyzed in each experiment. *n=* 7 independent experiments. Error bars indicate s.d. Statistical analysis was performed by unpaired two-tailed Student’s t-test. P-values are indicated. (d) Schematic of NE and DE differentiations for flow cytometry. (e) Flow cytometry quantification and representative histograms PAX6 Day 3 of NE differentiation using WT and *OCT4* KO hESCs. 2 WT and 2 KO lines analyzed per experiment. *n=* 3 independent experiments. Error bars indicate s.d. Statistical analysis was performed by unpaired two-tailed Student’s t-test. P-values are indicated. (f) Flow cytometry quantification and representative histograms SOX17 Day 3 of DE differentiation using WT and *OTUD5* KO hESCs. 2 WT and 2 KO lines analyzed per experiment. *n=* 3 independent experiments. Error bars indicate s.d. Statistical analysis was performed by unpaired two-tailed Student’s t-test. P-values are indicated.

